# Genetic modifiers of risk and age at onset in *GBA* associated Parkinson’s disease and Lewy body dementia

**DOI:** 10.1101/738351

**Authors:** Cornelis Blauwendraat, Xylena Reed, Lynne Krohn, Karl Heilbron, Sara Bandres-Ciga, Manuela Tan, Raphael Gibbs, Dena G. Hernandez, Ravindran Kumaran, Rebekah Langston, Luis Bonet Ponce, Roy N. Alcalay, Sharon Hassin-Baer, Lior Greenbaum, Hirotaka Iwaki, Hampton Leonard, Francis P. Grenn, Jennifer A. Ruskey, Marya Sabir, Sarah Ahmed, Mary B. Makarious, Lasse Pihlstrøm, Mathias Toft, Jacobus J. van Hilten, Johan Marinus, Claudia Schulte, Kathrin Brockmann, Manu Sharma, Ari Siitonen, Kari Majamaa, Johanna Eerola-Rautio, Pentti J. Tienari, The 23andMe Research Team, Alexander Pantelyat, Argye E. Hillis-Trupe, Ted M. Dawson, Liana S. Rosenthal, Marilyn S. Albert, Susan M. Resnick, Luigi Ferrucci, Christopher M. Morris, Olga Pletnikova, Juan Troncoso, Donald Grosset, Suzanne Lesage, Jean-Christophe Corvol, Alexis Brice, Alastair J. Noyce, Eliezer Masliah, Nick Wood, John Hardy, Lisa M. Shulman, Joseph Jankovic, Joshua M Shulman, Peter Heutink, Thomas Gasser, Paul Cannon, Sonja W. Scholz, Huw Morris, Mark Cookson, Mike A. Nalls, Ziv Gan-Or, Andrew B. Singleton, on behalf of the International Parkinson’s Disease Genomics Consortium (IPDGC)

**Author notes:** These authors have contributed equally (Ziv Gan-Or, Andrew B. Singleton). Correspondence to: Cornelis Blauwendraat, Laboratory of Neurogenetics, NIA, NIH, Building 35, 35 Convent Drive, Bethesda, MD 20892, USA.

## Abstract

Parkinson’s disease (PD) is a genetically complex disorder. Multiple genes have been shown to contribute to the risk of PD, and currently 90 independent risk variants have been identified by genome-wide association studies. Thus far, a number of genes (including *SNCA*, *LRRK2*, and *GBA*) have been shown to contain variability across a spectrum of frequency and effect, from rare, highly penetrant variants to common risk alleles with small effect sizes. Variants in *GBA*, encoding the enzyme glucocerebrosidase, are associated with Lewy body diseases such as PD and Lewy body dementia (LBD). These variants, which reduce or abolish enzymatic activity, confer a spectrum of disease risk, from 1.4- to >10-fold. An outstanding question in the field is what other genetic factors that influence *GBA*-associated risk for disease, and whether these overlap with known PD risk variants.

Using multiple, large case-control datasets, totalling 217,165 individuals (22,757 PD cases, 13,431 PD proxy cases, 622 LBD cases and 180,355 controls), we identified 1,772 PD cases, 711 proxy cases and 7,624 controls with a *GBA* variant (p.E326K, p.T369M or p.N370S). We performed a genome-wide association study and analysed the most recent PD-associated genetic risk score to detect genetic influences on *GBA* risk and age at onset. We attempted to replicate our findings in two independent datasets, including the personal genetics company 23andMe, Inc. and whole-genome sequencing data. Our analysis showed that the overall PD genetic risk score modifies risk for disease and decreases age at onset in carriers of *GBA* variants. Notably, this effect was consistent across all tested *GBA* risk variants. Dissecting this signal demonstrated that variants in close proximity to *SNCA* and *CTSB* (encoding cathepsin B) are the most significant contributors. Risk variants in the *CTSB* locus were identified to decrease mRNA expression of *CTSB*. Additional analyses suggest a possible genetic interaction between *GBA* and *CTSB* and *GBA* p.N370S neurons were shown to have decreased Cathepsin B expression compared to controls. These data provide a genetic basis for modification of *GBA*-associated PD risk and age at onset and demonstrate that variability at genes implicated in lysosomal function exerts the largest effect on *GBA* associated risk for disease. Further, these results have important implications for selection of *GBA* carriers for therapeutic interventions.

## Introduction

Heterozygous functional *GBA* variants are one of the most common genetic risk factors for Parkinson’s disease (PD) and Lewy body dementia (LBD), found in 3-20% of patients in different populations (Blauwendraat et al. 2018b; Gan-Or et al. 2015c; Guerreiro et al. 2018; Lesage et al. 2011; Rivas et al. 2018). In a homozygous or compound heterozygous state, *GBA* variants may cause Gaucher disease, an autosomal recessive lysosomal storage disorder. *GBA* encodes the lysosomal enzyme glucocerebrosidase (GCase), and it is hypothesized that loss of GCase activity leads to a reduced ability to degrade α-synuclein, encoded by *SNCA*. Aggregated, misfolded α-synuclein protein deposits in the form of Lewy bodies and Lewy neurites are the pathological hallmarks of PD and LBD.

Notably, *GBA* variants that do not cause Gaucher disease but do confer increased risk for PD and LBD have been identified. It is hypothesized that while these variants result in reduced GCase activity, the activity is not low enough to cause Gaucher disease. Multiple rare *GBA* variants have been described in PD in different populations. More common variants include p.E326K, p.T369M, p.N370S and p.L444P, whose frequencies vary with ethnicity and are each found on different haplotypes (Blauwendraat et al. 2018b; Leija-Salazar et al. 2019). In some ethnicities, such as the Ashkenazi Jewish population, certain *GBA* variants are found in about 5% of unaffected individuals, and 17-20% of PD patients (Gan-Or et al. 2008; Ruskey et al. 2019; Sidransky et al. 2009).

In European populations, the p.E326K risk variant is found in approximately 2% of the general population (https://gnomad.broadinstitute.org/). However, the majority of *GBA* variant carriers will not develop PD, implying that there are other genetic and/or environmental factors that affect the penetrance of these variants. Studies that have examined the penetrance of *GBA* variants in carriers suggest it is age-related and is typically between 10-30% (Anheim et al. 2012; Rana et al. 2013). Furthermore, penetrance differs across *GBA* variants, with high-risk variants leading to earlier disease onset compared to lower risk variants (Gan-Or et al. 2015a). Interestingly, it was demonstrated that various *GBA* variants, including p.E326K, p.T369M and p.N370S have similar effects on GCase activity in humans, reducing it by 18-46% on average (Alcalay et al. 2015). Additionally, genotype-phenotype studies have shown that PD patients with *GBA* variants have an earlier age at onset (AAO), faster disease progression, and higher rates of non-motor symptoms, such as rapid eye movement (REM) sleep behaviour disorder (RBD), autonomic dysfunction, hallucinations and cognitive decline, compared to those with non-*GBA* associated PD (Gan-Or et al. 2018).

In recent years, *GBA* has become a prominent target for therapeutic development, and the first gene-specific phase 2 clinical trial in PD is currently ongoing for *GBA*-positive PD patients (ClinicalTrials.gov Identifier: NCT02906020). One of the concerns in performing such a trial is that, despite the randomization for treatment and placebo groups, these groups will remain unbalanced in terms of factors that affect their progression, which can be a significant confounder of trial outcome (Leonard et al. 2018). In addition, it is likely that future preventive clinical trials will also target populations that are at high risk for Lewy body diseases, such as *GBA* risk variant carriers with prodromal symptoms. Therefore, identifying factors that can affect the penetrance and clinical presentation of *GBA*-associated PD will be crucial for trial design. Furthermore, modifiers of penetrance AAO may also become drug targets.

In the current study, we gathered data from multiple sources, including the International Parkinson’s Disease Genomics Consortium (IPDGC) and the UK Biobank (UKB), to identify *GBA* variants in both cases and controls and used 23andMe and whole-genome sequencing data for further validation. Subsequently, we used genome-wide association studies (GWAS) and genetic risk scoring to identify genetic variants that modify the penetrance and AAO of *GBA*-associated PD. We then examined protein levels of the top hits in forebrain neurons differentiated from iPS cells of individuals with and without *GBA* variants.

## Materials and methods

### Genotyping data

#### International Parkinson Disease Genomics Consortium genotyping data

Genotyping data (all Illumina platform based) was obtained from International Parkinson’s Disease Genomics Consortium (IPDGC) members, collaborators, and public resources (Supplementary Table 1 and 2). All datasets underwent quality control separately, both on individual-level data and variant-level data before imputation as previously described (Blauwendraat et al. 2019; Nalls et al. 2018). Refer to Supplementary data for a detailed description of data processing.

For GWAS analyses, variants passing the following post-imputation criteria were included: R^2^ > 0.8, minor allele frequency > 0.05, and genotype missingness < 0.15 across subsets. Case-control GWAS were performed using RVTESTS with default settings (Zhan et al. 2016) using logistic regression with sex, age (if not available the mean age of all data was used), principal components (PC) 1-5, and dataset origin as covariates. AAO GWAS was performed in *GBA* carriers only using RVTESTS linear regression with AAO as a continuous phenotype and sex, PC 1-5 and dataset origin as covariates. Cases without age information were excluded from the AAO GWAS, and individuals with two *GBA* variants were excluded from all analyses to prevent bias (Supplementary Table 2).

#### Lewy body dementia genotyping data

LBD cases and controls were genotyped for ongoing projects at the Neurodegenerative Diseases Research Unit (NDRU) using the NeuroChip genotyping array (Illumina, San Diego, CA, USA). Genotyping was performed as previously described (Blauwendraat et al. 2017). All cases met the McKeith criteria for probable or definite disease (McKeith 2006). For more details on this dataset, see Supplementary Table 2. Genotype quality control, processing, and analysis was performed similar to the above described IPDGC genotype data. For analyses, variants that passed the following post-imputation criteria were included: imputation R^2^ > 0.8 and minor allele frequency > 0.05. Case-control analyses were performed using RVTESTS using logistic regression on genotype dosages with sex, and PC 1-5 as covariates. Age was not used since the majority of the individuals had only age of death information available.

#### McGill GBA genotyping data

Ashkenazi Jewish PD cases were genotyped at McGill University using the Illumina Human OmniExpress Array + custom SNPs of the NeuroX array (Nalls et al. 2015b). Sanger sequencing or targeted next-generation sequencing confirmed *GBA* p.N370S status. For more details on this dataset, see Supplementary Table 2. *GBA* variant carriers from the McGill dataset included samples from Columbia University, NY, USA (Alcalay et al. 2015) and Sheba Medical Center, Ramat Gan, Israel (Ruskey et al. 2019). Genotype quality control, processing, and analysis was performed similar to the above described genotyping data.

#### UK Biobank data

Imputed UKB genotype data (v3) was downloaded (April 2018) under application number 32663 (Bycroft et al. 2018; Sudlow et al. 2015). All European ancestry individuals (based on code 22006) were screened for the common *GBA* coding variants p.E326K, p.T369M (both directly genotyped) and p.N370S (imputed R^2^=0.81). PD cases were identified using ICD10 code “G20” from data fields 41202 (Diagnoses - main ICD10), 41204 (Diagnoses - secondary ICD10) and 40002 (Contributory (secondary) causes of death: ICD10). Additionally, self-report PD cases were included from data field 20002 with code “1262”. “Proxy” PD cases were included by using individuals who reported a parent with PD (paternal PD, data field 20107 and maternal PD, data field 20110) which have been previously shown to share genetic risk with PD cases (Nalls et al. 2018). Controls were set as people with no report of PD, no parent with PD and no sibling with PD (data field 20111). Covariates were obtained from the data fields: genetic sex (22001) batch (22000), age of recruitment (21022), and Townsend index (189). Individuals were filtered for relatedness (PIHAT>0.125) based on the pre-imputed data. Imputed genotypes were converted to PLINK2 .pgen files using PLINK2 (version v2.00) and filtered for missingness (<0.05) and minor allele frequency (>0.05), Hardy-Weinberg equilibrium of P ≥1E-6 in controls and imputation quality (R^2^ > 0.8). GWAS was performed using PLINK2 logistic regression with covariates including batch, age of recruitment, Townsend index, and 5 PCs created using FlashPCA (Abraham et al. 2017). All UKB analyses were performed using population controls with age of recruitment ≥60 and for *GBA* carrier case-control analyses, the case-control ratio was set to 1:9 by randomly selected controls.

#### Additional datasets

Additional results for variants of interest were obtained from 23andMe and whole-genome sequencing data obtained from various resources. Analyses were performed using similar and identical methods as described above. Associations in 23andMe were performed using logistic regression correcting for age, sex, genotyping platform (v4 or v5) and 5 PCs. Genome-sequencing data was obtained from multiple ongoing projects at the Laboratory of Neurogenetics, see Supplemental Data for a detailed description. Sample sizes can be found in Supplementary Table 3.

### Genetic risk score analyses

To assess the influence of the genetic risk score (GRS), we calculated the GRS using the effect sizes from the most recent PD GWAS variants (Nalls et al. 2018) (see Supplementary Table 4). Note that most of the data are included in the latest PD GWAS, however, for the calculation of the GRS in the UKB data we excluded the UKB summary statistics. Overall, at least 65 of these variants of interest were imputed with high quality (R^2^ > 0.8) in the included datasets, allowing for a maximum of 15% variant missingness across any constituent data subset due to differing genotyping backbones. For the IPDGC dataset, rs356182 was replaced by rs356219, due to a higher imputation quality (R^2^ > 0.82 and D’ 0.959 between variants in the IPDGC reference panel). Variants from the *GBA* locus and *LRRK2* p.G2019 were excluded from the GRS in analyses including *GBA* carriers only. The GRS was calculated and processed using PLINK (v1.90) for each individual as described previously (Nalls et al. 2015a). To make the scale of analyses relating to the PD-GRS more easily interpretable, PD-GRS were converted to Z-scores. GRS associations were performed with a logistic regression in R (v3.5.1) using case-control status or linear regression with AAO as the outcome phenotype. As covariates, we included: sex, age (in case-control status comparison only), the first 10 PCs, and the dataset origin (due to the fact that the IPDGC data is derived from multiple sources see Supplementary Table 2). Area under the curve risk predictions based on the GRS were performed using the pROC package (Robin et al. 2011).

### Additional analyses

*GBA* variants were annotated using ANNOVAR (Wang et al. 2010). Nucleotide positions in *GBA* were based on build hg19 and amino acid positions were based on NM_001005742. Note that historically these *GBA* variants are also known as p.E365K → p.E326K (rs2230288), p.T408M → p.T369M (rs75548401), and p.N409S → p.N370S (rs76763715). Where possible, association results were meta-analysed with METAL (v.2011-03-25) using default settings (Willer et al. 2010) or using the metaphor R package (2.0) using fixed effects models (Viechtbauer 2010). We excluded SNPs with a heterogeneity I^2^ statistic > 80% and variants that were present in only one GWAS. Limited population stratification was seen despite low sample numbers in some GWAS (Supplementary Table 5). Differences in AAO between *GBA* variants was tested using linear regression in R using AAO as the dependent variable, correcting for sex, PCs 1-5, and the dataset origin as covariates. Conditional GCTA-cojo analyses (Jian Yang et al. 2011; J. Yang et al. 2012) were performed to identify whether there were multiple independent signals in loci of interest, using the previously described IPDGC reference panel (Blauwendraat et al. 2019; Nalls et al. 2018). Genetic interaction analyses were performed using R (version 3.5) using an interaction term created using SNP1+1 * SNP2+1, followed by a regression using both variants and covariates (PC1-5, sex and age if available).

Exploratory gene expression values and functional annotation of identified risk variants were obtained by mining datasets including: the largest brain eQTL study to date (Qi et al. 2018), the GTEx QTL portal V7 (https://gtexportal.org/) (Consortium 2013) and the NABEC dataset (phs001300.v1.p1, (Gibbs et al. 2010)). LocusZoom (Pruim et al. 2010) and LocusCompare plots (B. Liu et al. 2019) were generated for the regions of interest and were compared to the latest published PD GWAS. Single-nuclei RNAseq expression of three frozen prefrontal cortex samples and four frozen hippocampal samples were obtained from dbGaP accession phs000424.v8.p1 (Habib et al. 2017). Data was processed using Seurat and genes of interest were highlighted to explore expression patterns (Butler et al. 2018).

### *GBA* variant validation

To validate the genotyping and imputation of *GBA* variants, we obtained next-generation sequencing (targeted resequencing and genome sequencing) data from two parallel IPDGC projects. *GBA* carrier genotypes were merged using PLINK (v1.90) with both the resequencing and whole genome sequencing data. Overlapping individuals were identified using > 0.9 PIHAT values. Additionally, the majority of the PROBAND dataset individuals were Sanger sequenced for the full *GBA* gene as described elsewhere (Malek et al. 2018). In the McGill dataset, *GBA* variants were confirmed using Sanger sequencing or targeted next generation sequencing as previously described (Ruskey et al. 2019). Overall, the concordance rate between the imputed/genotype data and the sequence data for the presence of at least one p.N370S and p.T369M allele was 100% and for p.E326K was 98% and 94% in the next-generation and Sanger sequencing data, respectively (see Supplementary data for more details).

### Induced pluripotent stem cells and neuron analysis

#### Forebrain neuron differentiation

Induced pluripotent stem cells (iPSC) from the Parkinson’s Progression Marker Initiative (PPMI) were selected based on their genotype at *GBA* and rs1293298 then differentiated to forebrain type neurons. iPSCs were grown in E8 media (ThermoFisher) on Matrigel until 90% confluent. When cells grew to confluence they were transitioned to N3 media (50% DMEM/F12, 50% Neurobasal with 1x Penicillin-Streptomycin, 0.5x B-27 minus vitamin A, 0.5x N2 supplement, 1x Glutamax, 1x NEAA, 0.055mM 2-mercaptoethanol and 1ug/ml Insulin) plus 1.5uM Dorsomorphin (Tocris Bioscience) and 10uM SB431542 (Stemgent). Media was replaced every day for 11 days. On day 12 Dorsomorphin and SB431542 were removed and cells continued to be fed each day with N3. N3 was supplemented with 0.05uM Retinoic acid from days 16 through 20. The differentiating cells were split 1:2 with Accutase and seeded with ROCK inhibitor onto Poly-L-ornithine (Sigma), fibronectin (2ug/ml) and laminin (0.2ug/ml) coated plates on day 20. Cells were then fed daily with N4 media (same as N3 plus 0.05uM Retinoic acid, 2ng/ml BDNF and 2ng/ml GDNF) until day 26 when they were frozen in synthafreeze. Vials were thawed as needed in N4 media. Differentiation state was confirmed by immunocytochemistry for neuronal markers Beta-III Tubulin (Novus Biologicals, NB100-1612) and MAP2 (Santa Cruz Biotechnology, sc-20172) with nuclei counterstained for Hoechst 33342 (ThermoFisher Scientific, H3570).

#### Protein analysis

Protein lysates were collected in buffer with 20 mM TrisHCl pH 7.5, 150mM NaCl, 1mM EDTA, 0.3% Triton 10% Glycerol, 1x protease inhibitors, and 1x phosphatase inhibitors and run as previously reported (Blauwendraat et al. 2018a) on 4-20% Criterion TGX gels (Bio-Rad) in Tris-glycine running buffer and transferred to nitrocellulose membranes (Bio-Rad) using the Trans-blot Turbo system (Bio-Rad). Membranes were blocked in Odyssey Blocking buffer (LI-COR) diluted by 50% in PBS. Primary antibodies for Cathepsin B (Abcam, ab92955) and α-synuclein (BD Biosciences, 610787) were incubated overnight at 4 degrees. Primary antibody to β-III Tubulin (Novus Biologicals, NB100-1612) was incubated for one hour at room temperature. LI-COR secondary IRDye antibodies were incubated 1 hour at room temperature. Proteins were visualized on the Odyssey CLx Imaging System (Licor) and quantified using Image Studio Lite (Licor).

## Results

### Initial data overview

In total we included 21,478 PD cases and 24,388 controls from IPDGC datasets; 1,176 PD cases, 13,431 proxy PD cases, and 155,325 population controls from the UKB dataset; 103 *GBA* p.N370S PD cases from the McGill dataset; and 622 LBD cases and 782 controls from the NDRU dataset (Supplementary Table 2). In the HRC reference panel, 10 coding *GBA* variants are present (Supplementary Table 6), with varying imputation qualities since common variants are more easily imputed than variants with lower frequencies. Additionally, some *GBA* variants were directly genotyped in some datasets (Supplementary Table 1). The three most common variants (p.E326K, p.T369M and p.N370S) had the highest quality, we therefore selected these three variants for all subsequent analyses. This resulted in a total of 1,772 *GBA* carriers with PD, 81 with LBD, 711 PD proxy cases, and 7,624 control *GBA* carriers. We excluded individuals carrying two *GBA* variants to ensure conservative estimates and reduce bias due to the association with Gaucher disease (Supplementary Table 2). All included *GBA* variants (p.E326K, p.T369M and p.N370S) have previously been shown to be enriched in cases in all datasets.

### Parkinson’s disease genetic risk score modifies disease risk in *GBA* carriers

Exploring the potential effect of PD case-control GWAS loci, we calculated the PD-GRS, both including and excluding *GBA* variants and *LRRK2* p.G2019S for all three datasets. To compare the effect size associated with the GRS, we examined four groups within each dataset: controls without a *GBA* risk variant, PD cases without a *GBA* risk variant, controls with a *GBA* risk variant, and PD cases with a *GBA* risk variant. Additionally, for the UKB dataset, we included proxy-PD cases (having a parent with PD) (Figure 1). As expected, the GRS is significantly associated with risk in *GBA*-negative PD cases compared to controls and interestingly, also in the NDRU-LBD dataset (Figure 1 and Table 1). This effect is highly similar between *GBA* carrier controls and *GBA* carrier cases, when excluding *GBA* and *LRRK2* variants from the genetic risk score (Figure 2 and Table 1). This effect is consistent in subsets based on all three *GBA* variants (Supplementary Figure 1). Although this was not significant in the (smaller) p.N370S group, the direction and effect size were highly similar to the other *GBA* variants (Supplementary Figure 1, ranging ORs from 1.31 to 1.55). Notably, we do not observe a consistent difference in genetic risk between PD or LBD cases with and without *GBA* variants when excluding *GBA* and *LRRK2* in the genetic risk (Table 1).

**Figure 1:**
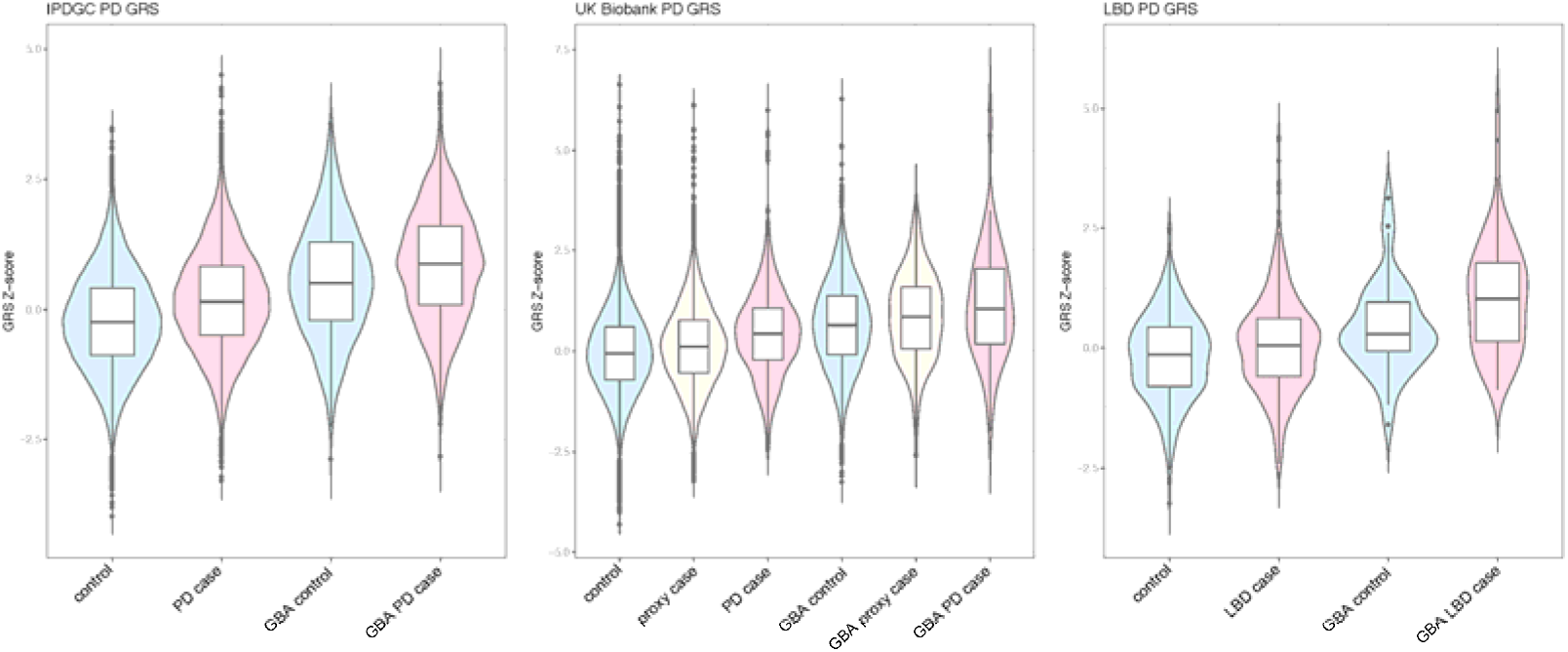
Parkinson disease genetic risk score boxplots divided by disease status and *GBA* carrier status. Genetic risk score differences between *GBA* negative controls, *GBA* negative Parkinson’s disease or Lewy body dementia cases, *GBA* positive controls and *GBA* positive cases. In each dataset, a highly similar effect is observed clearly separating the subgroups.

**Table 1:** Associations between the Parkinson disease genetic risk score and case-control status. Genetic risk score associations performed per group. ORs are based on a Z-score scale as described in the methods. UKB= UK Biobank, SE=standard error * For these associations, the *GBA* region and *LRRK2* p.G2019S were excluded when calculating the genetic risk score.

| Outcome | Case-control |  | GBA+, Case-control* |  | GBA+ case vs GBA- case* |  |
| --- | --- | --- | --- | --- | --- | --- |
|  | P | OR (95%CI) | P | OR (95%CI) | P | OR (95%CI) |
| IPDGC | 1.877E-267 | 1.531 (1.51-1.56) | 2.75E-12 | 1.464 (1.36-1.57) | 0.00747 | 0.926 (0.87-0.98) |
| UKB | 9.156E-55 | 1.557 (1.5-1.61) | 1.96E-4 | 1.578 (1.34-1.82) | 0.674 | 1.050 (0.82-1.28) |
| UKB proxy-case | 1.284E-35 | 1.138 (1.12-1.16) | 2.12E-5 | 1.217 (1.13-1.31) | NA | NA |
| NDRU-LBD | 6.88E-8 | 1.370 (1.26-1.48) | 0.0798 | 1.565 (1.06-2.07) | 0.892 | 1.017 (0.78-1.26) |

**Figure 2:**
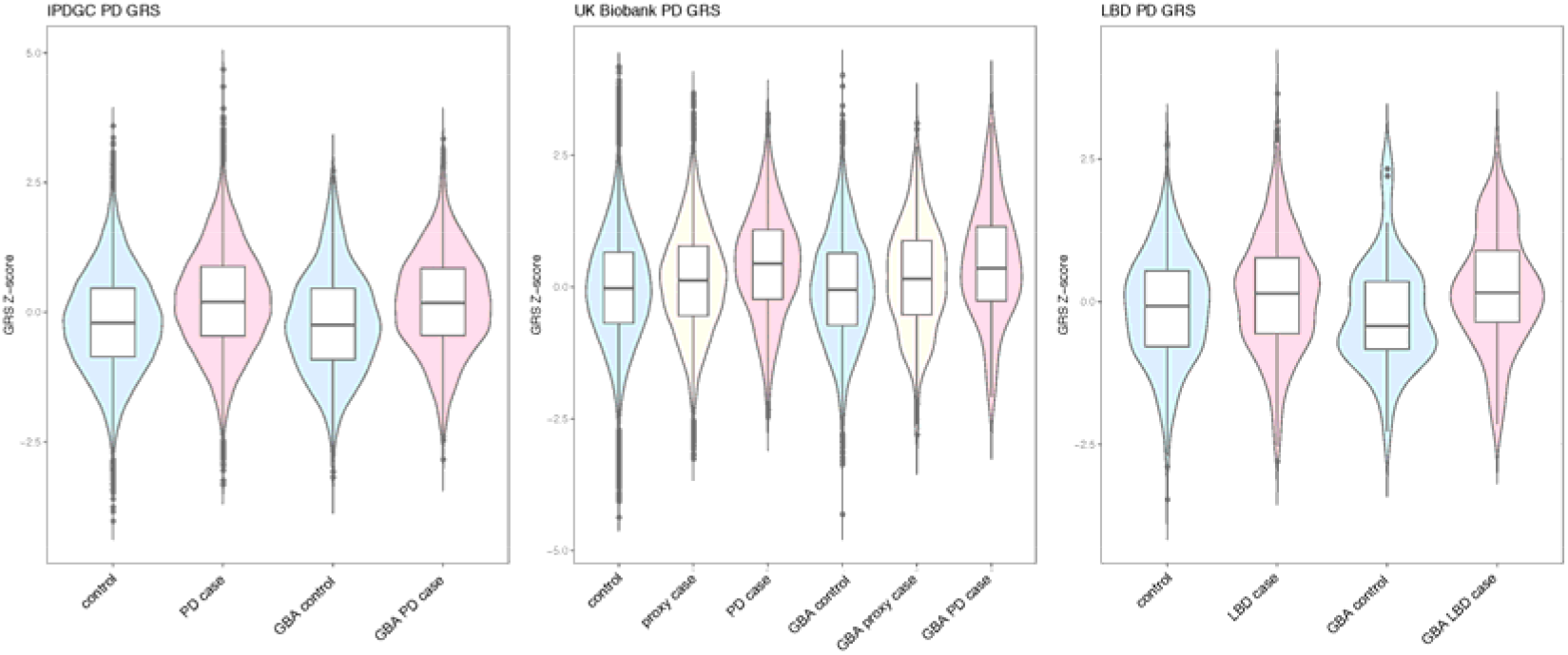
Parkinson disease genetic risk score boxplots divided by disease status and *GBA* carrier status excluding *GBA* in the genetic risk score calculation. Genetic risk score differences between *GBA* negative controls, *GBA* negative Parkinson’s disease or Lewy body dementia cases, *GBA* positive controls and *GBA* positive cases when excluding the *GBA* region and *LRRK2* p.G2019S from the calculations. In each dataset, a highly similar effect is observed clearly separating the cases and controls. However, there is no clear separation between *GBA* cases and *GBA* negative cases or *GBA* negative controls and *GBA* positive controls.

When assessing the predictive power of the GRS in *GBA* positive cases versus *GBA* positive controls (excluding *GBA* and *LRRK2* variants in GRS calculations), these values were slightly lower compared to the reported predictive power of the GRS in PD cases as previously described by Nalls et al (Nalls et al. 2018) (AUC=0.651, 95%CI 0.617-0.684), with AUC of 0.614 (95%CI 0.591-0.637) in the IPDGC dataset and 0.622 (95%CI 0.561-0.684) in the UKB dataset (Supplementary Table 7 and Supplementary Figure 2-4). As expected, the NDRU-LBD dataset performed poorer than the PD cases with an AUC of 0.570 (95%CI 0.540-0.600), supporting the notion of partial overlapping and shared genetic risk between PD and LBD.

### *SNCA* and *CTSB* loci as genetic modifiers of disease risk in *GBA* carriers

To identify potential genetic modifiers of *GBA*-associated PD penetrance, we performed a GWAS comparing PD patients with *GBA* variants and controls with *GBA* variants. We first examined the association using all three *GBA* variants together (p.E326K, p.T369M and p.N370S) in the IPDGC and the UKB dataset, followed by a meta-analysis of both. After meta-analysing the *GBA* data in both datasets, no genome-wide significant variants were identified (Supplemental Figure 5).

When solely examining the 90 PD variants previously identified in a PD case-control GWAS (Nalls et al. 2018), variants in two PD loci passed Bonferroni multiple test correction for association with PD, *SNCA* and *CTSB*. The strongest association was observed with rs356219 in the *SNCA* locus (meta-P=9.26E-07, OR=1.375, CI95%=1.211-1.563, Table 2 and Supplemental Figure 6). In addition to *SNCA*, the *CTSB* locus rs1293298 (meta-P=9.74E-06, OR=0.725, CI95%=0.629-0.836, Table 2 and Supplemental Figure 7) was also significantly associated with *GBA* penetrance after correction for multiple testing (Supplementary Table 8). Additional investigation of these two variants was performed in the 23andMe PD case-control dataset and in a WGS dataset (Table 2). Meta-analyses of these results showed a genome-wide significant P-value (P=2.23E-11) for the *SNCA* variant and a more significant P-value for the *CTSB* variant (P=6.34E-6) (Table 2 and Figure 3). The majority of these effects are mostly driven by p.E326K carriers likely due to the larger sample size, but the effect size was consistent for each of the three variants when tested individually in the IPDGC dataset (Supplementary Table 9, Supplementary Figure 6 and 7).

**Table 2:** Meta-analyses of significant GWAS signals and interaction effects. Two variants were significant after correction of multiple testing: rs356219 (*SNCA*) and rs1293298 (*CTSB*) in the initial used datasets. Additional datasets were assessed and overall both P-values became more significant. The interaction effects between the two variants and *GBA* was not replicated in the external datasets.

| Association | IPDGC (P-value) | IPDGC (OR, CI) | UKB (P-value) | UKB (OR, CI) | Meta (P-value) | Meta (OR, CI) | 23andMe (P-value) | 23andMe (OR, CI) | LNG Genomes (P-value) | LNG Genomes (OR, CI) | Meta_all (P-value) | Meta_all (OR, CI) |
| --- | --- | --- | --- | --- | --- | --- | --- | --- | --- | --- | --- | --- |
| rs356219 (*SNCA*) | 6.37E-05 | 1.315 (1.150-1.504) | 3.99E-04 | 2.085 (1.388-3.132) | 9.26E-07 | 1.376 (1.211-1.563) | 6.89E-07 | 1.273 (1.157-1.400) | 0.6645 | 0.921 (0.635-1.336) | 2.23E-11 | 1.290 (1.198-1.391) |
| rs1293298 (*CTSB*) | 8.27E-05 | 0.740 (0.637-0.860) | 0.0304478 | 0.614 (0.394-0.955) | 9.74E-06 | 0.726 (0.630-0.837) | 0.0564 | 0.898 (0.805-1.003) | 0.05908 | 0.6616 (0.431-1.016) | 6.34E-06 | 0.822 (0.755-0.895) |
| **Interaction** |  |  |  |  |  |  |  |  |  |  |  |  |
| rs356219 * GBA | 0.7025 | 1.029 (0.889-1.190) | 0.00639 | 1.600 (1.141-2.244) | 0.152 | 1.103 (0.965-1.261) | 0.773 | 0.985 (0.891-1.089) | 0.3697 | 0.823 (0.537-1.261) | 0.653 | 1.018 (0.941-1.102) |
| rs1293298 * GBA | 0.00647 | 0.796 (0.676-0.938) | 0.28833 | 0.802 (0.534-1.205) | 0.00347 | 0.797 (0.685-0.928) | 0.554 | 0.966 (0.860-1.084) | 0.0663 | 0.625 (0.379-1.032) | 0.0113 | 0.890 (0.813-0.974) |

**Figure 3:**
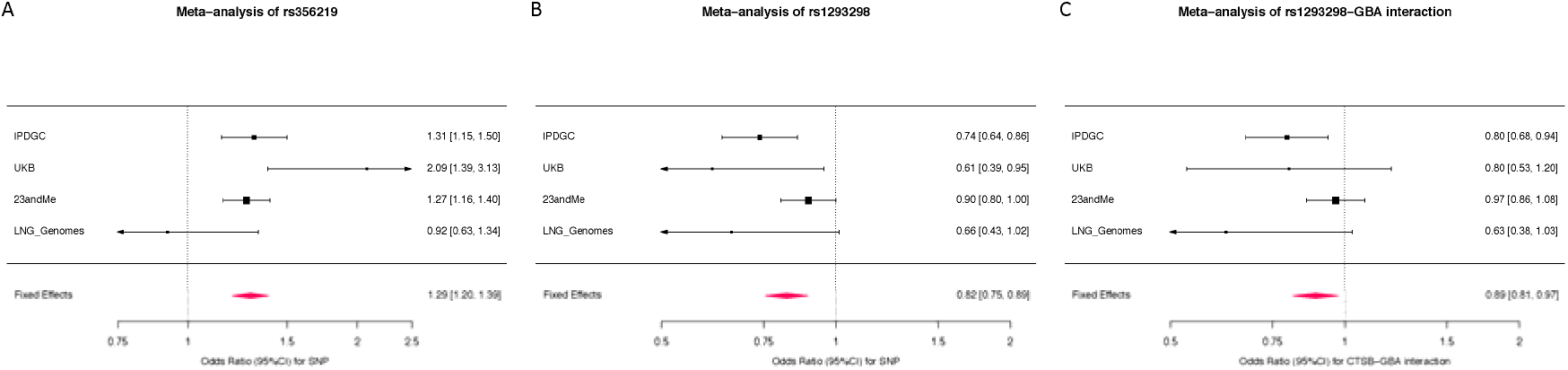
Meta-analyses of significant GWAS signals and interaction effect. A) Meta-analysis of the rs356219 variant in all four datasets resulted in a significant effect P=2.23E-11 and I2=65.4. B) Meta-analysis of the rs1293298 variant in all four datasets resulted in a significant effect P=6.34E-6 and I2=57.4. C) Meta-analysis of the interaction between the rs1293298 variant and having a damaging *GBA* variant resulted in a significant effect P=6.34E-6 and I2=48.8.

### Evidence for a potential genetic interaction between *CTSB* and *GBA*

Interestingly, the OR of the *CTSB* variant rs1293298 association in the *GBA* GWAS is about 3-fold lower compared to the PD case-control GWAS (Table 2, (Nalls et al. 2018) OR =0.911, CI95%=0.891-0.932 (Supplemental Figure 8), while for the *SNCA* variant this was not the case. This suggests a potential interaction between *CTSB* and *GBA*. Examining this possibility further resulted in a significant interaction term after meta-analysing effects of the IPDGC and UKB data (Table 2). As expected, no consistent significant interaction term was identified between *SNCA* and *GBA* (Table 2, Supplemental Figure 9). Replication of these results was investigated in the 23andMe and LNG Genomes data for both interactions. In both datasets the interaction terms were not significant, although the direction of effect was the same for *CTSB* x *GBA* (Table 2 and Figure 3).

### Genetic dissection of the *CTSB* GWAS locus

To evaluate the potential molecular mechanism for increased risk of PD in *GBA* carriers, we explored gene expression and eQTL datasets for the two significant associated loci, *SNCA* (rs356219) and *CTSB* (rs1293298). For the *SNCA* locus, several previous reports showed that risk variants in this locus likely increase *SNCA* expression (Pihlstrøm et al. 2018; Soldner et al. 2016). For the *CTSB*, no comprehensive assessment has yet been performed. In both the PD GWAS and the *GBA* GWAS, rs1293298 appears to be the index variant and both GWAS signals appears to be highly similar (Figure 4 A,B,C).

**Figure 4:**
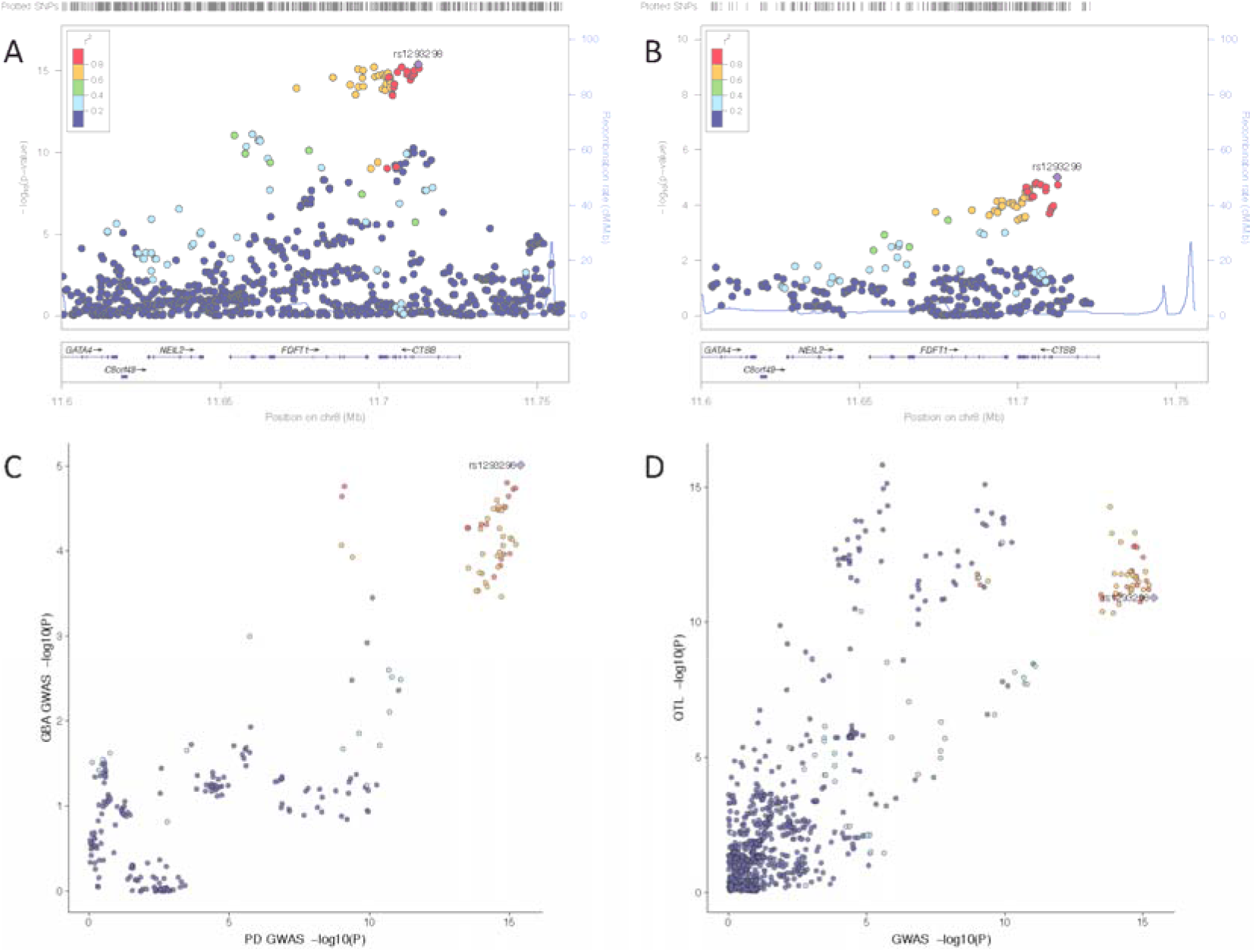
Genetic dissection of the *CTSB* locus. A) LocusZoom plot of the *CTSB* region from the latest PD GWAS (Nalls et al 2019). B) LocusZoom plot of the *CTSB* region from the latest current *GBA* carrier GWAS. C) LocusCompare plot of the *CTSB* region showing high correlation between the PD GWAS (x-axis) and the *GBA* carrier GWAS (y-axis). D) LocusCompare plot of the *CTSB* region showing high correlation between the PD GWAS (x-axis) and the brain QTL data (Qi et al 2019) (y-axis).

The protective allele (A) of the *CTSB* variant rs1293298 increases *CTSB* expression in the brain in both GTEx and NABEC datasets and was significant in the latest meta-analyses of brain eQTL data (Supplementary Table 10). Results from the NABEC dataset suggest that specific isoforms of *CTSB* are differentially regulated and the QTL effect is mainly for the most highly expressed transcript (isoform: uc003wuq.3, Supplementary Figure 10). Interestingly the QTL signal shows high correlation with the PD GWAS signal (Figure 4D). Exploring post-mortem brain-derived single nuclei RNAseq data showed that *CTSB* is mainly expressed in neurons and microglia in a pattern that is similar to *SNCA* expression (Supplementary Figure 11). This suggests that the biological effects of these variants possibly takes place in these cell types, which is in line with pathology and previous literature. We attempted to examine *GBA* expression levels, but they were too low to assess.

Given the function of Cathepsin B (encoded by *CTSB*) as lysosomal protease, which is of interest for PD pathogenesis in general (McGlinchey and Lee 2015), we explored other members of the cathepsin family for: 1) genetic association in the most recent PD GWAS; 2) genetic association in *GBA* GWAS and 3) the presence of an eQTL at each locus. In addition to the peak at *CTSB*, only *CTSE* has a PD GWAS peak in close proximity, however this locus is already nominated as the *NUCKS1*/*RAB29* locus (Supplementary Table 11). Interestingly, for almost all cathepsins a brain eQTL has been identified. However, with the exception of *CTSB*, none of the eQTLs were associated with PD (Supplementary Table 11). These results suggest that expression differences in other cathepsins, including *CTSD* and *CTSL* which were previously suggested to degrade alpha-synuclein (McGlinchey and Lee 2015), do not modulate risk for disease.

### Cathepsin B and α-synuclein levels in iPSC-derived neurons from GBA carriers and non-carriers

We examined the potential relationship between *GBA* and *CTSB* using PPMI iPSCs differentiated to forebrain neurons. For this analysis we used iPSCs from individuals with and without the *GBA* p.N370S variant to interrogate Cathepsin B and α-synuclein protein levels in the context of *GBA* variants. All lines used were heterozygous for rs1293298 at the *CTSB* locus and neuronal differentiation was confirmed by immunocytochemistry for β-III tubulin and MAP2 (Supplementary figure 12). We observed a significantly lower (P = 0.0291; unpaired t-test, n=6) level of processed Cathepsin B protein in *GBA* variant carriers compared to cells without *GBA* variants (*GBA* WT = 0.528 ± 0.0930; *GBA* p.N370S = 0.263 ± 0.0467) (Figure 5). These results show that the *GBA* variant carriers have reduced levels of mature Cathepsin B which may result in even lower lysosomal protease activity and increased accumulation of protein aggregates in neurons. Although the forebrain neurons carrying the *GBA* variant did not have increased levels of α-synuclein (Supplementary Figure 13) it is possible that the overall reduction in lysosomal proteases allows for a faster accumulation of α-synuclein aggregates as neurons age.

**Figure 5:**
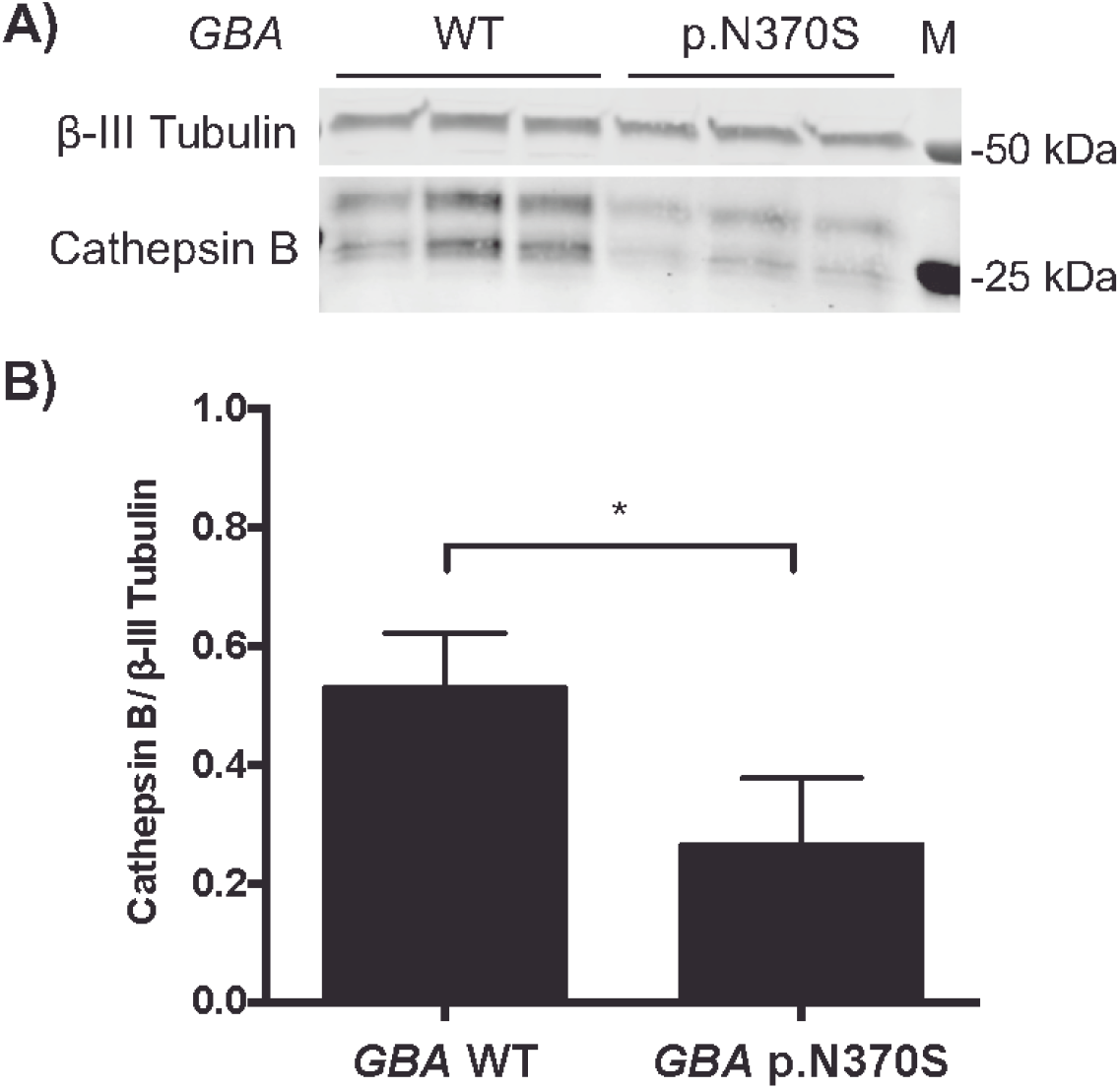
Cathepsin B protein expression is decreased in iPSC derived neurons from *GBA* p.N370S carriers. Forebrain neurons differentiated from patient iPS cells with *GBA* mutations have decreased Cathepsin B protein expression. A) Representative western blot showing protein levels of processed Cathepsin B (CTSB, 24 and 27 kDa) and neuron associated β-III Tubulin (TUBB3, 50 kDa) in forebrain neurons differentiated from individuals with and without PD-associated *GBA* variants (n=2 lines from each genotype). B) Quantification of the mean processed Cathepsin B protein levels relative to TUBB3 shows a significant decrease (* P<0.05; unpaired t-test, n=2 lines of each genotype with 3 technical replicates each) in Cathepsin B levels in forebrain neurons differentiated from *GBA* p.N370S carriers compared to neurons from WT *GBA* carriers. M = protein marker; error bars represent standard error of the mean.

### Genetic modifiers of age at onset in *GBA* associated Parkinson’s disease cases

Information on AAO was available for the majority of the IPDGC *GBA* PD cases (89.9% n=1353 cases, average=60.54 years, SD=11.42). Cases with two *GBA* variants (either homozygous or compound heterozygous, n=38) had a lower AAO compared to single variant carriers; however, no statistically significant difference was detected (P= 0.107; linear regression, Beta=−2.93 years, SE=1.82, two *GBA* variant case average=56.81, SD=10.50 vs single *GBA* variant case average=60.54, SD=11.42). The AAO of *GBA* PD cases is significantly lower compared to non-*GBA* PD cases (linear regression, P=9.70E-06, Beta=−1.43, SE=0.324, *GBA* PD case average=56.81, SD=10.50 vs non-*GBA* case average=60.54, SD=11.42). On average p.N370S cases had the earliest AAO followed by p.E326K and p.T369M (Figure 6C). Of note, the *LRRK2* p.G2019S variant was also identified in 14 *GBA* carriers (9 p.N370S and 5 p.E326K carriers) and did not seem to influence the AAO P=0.44 (Linear regression, Beta=2.41, SE=3.15, GBA-PD-*LRRK2* p.G2019S carriers average=63.86, SD=10.87 vs *GBA*-PD-non-*LRRK2* p.G2019S average=60.29, SD=11.80), which is consistent with previous findings, although power was limited given the small number of included individuals (Yahalom et al. 2018).

**Figure 6:**
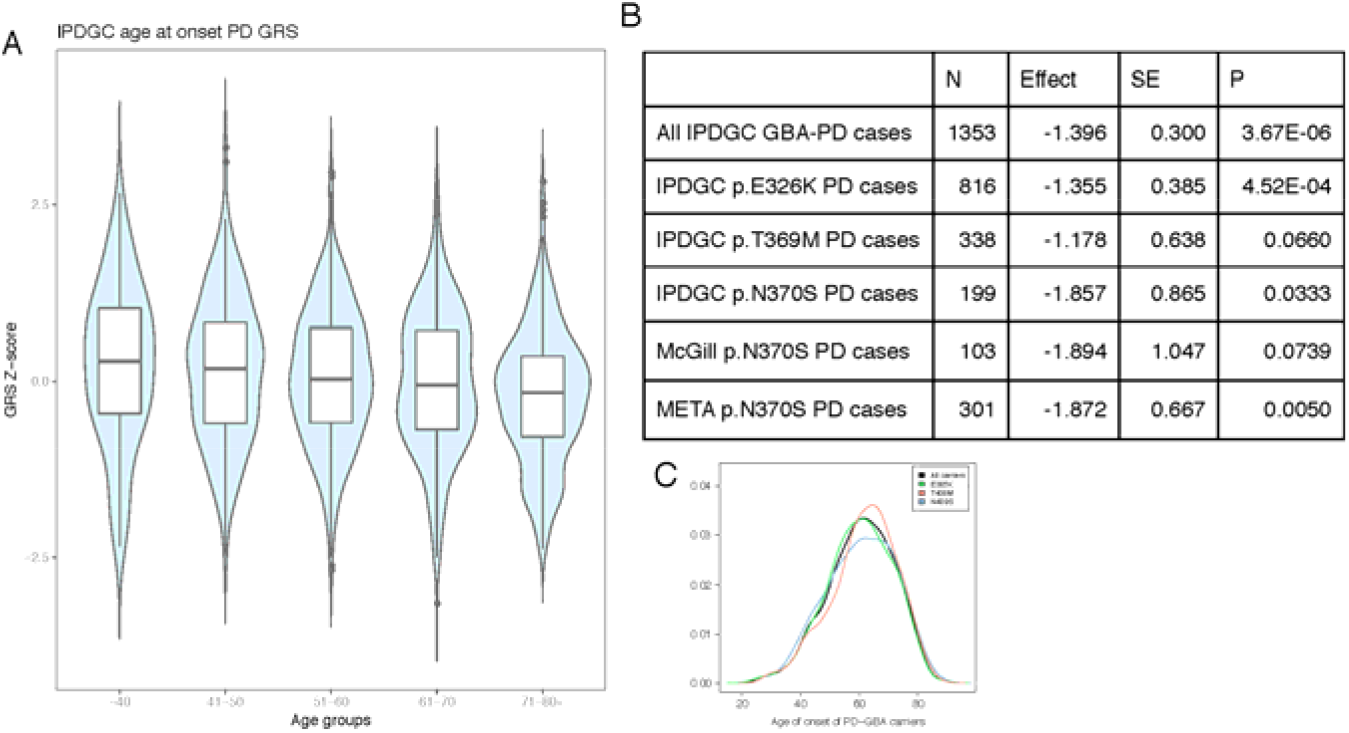
Parkinson’s disease genetic risk score significantly influences the age at onset of *GBA* positive cases. A) Parkinson’s disease cases carrying a *GBA* variant were grouped by ages and a clear and significant decline was seen for the genetic risk score. B) The effect size was similar across all tested *GBA* variants (estimates relate to the rare allele). C) When plotting the AAO distribution of all tested *GBA* variants, p.N370S Parkinson’s disease cases had an earlier AAO followed by p.E326K and p.T369M.

To identify potential *GBA* AAO modifiers, we performed a GWAS using AAO as phenotype in all *GBA* positive PD cases (n=1,353). No genome-wide hits were identified, most likely due to the relatively low sample size. When investigating previously identified PD loci, no variants passed Bonferroni correction. However, nominally significant signals were identified for several PD loci, including *SNCA* and *TMEM175* (Supplementary Table 12). Similar to PD in general, the PD-GRS is significantly associated with a lower AAO in *GBA* carriers (Figure 6A) (Blauwendraat et al. 2019). This effect was mainly driven by p.E326K carriers. However, both p.T369M and p.N370S carriers also show the same trend towards lower AAO with a similar effect size that is close to significance likely due to lower sample numbers (Figure 6B). This association was similar in an independent p.N370S Ashkenazi Jewish ancestry dataset, again demonstrating the same general trend and similar effect size. A meta-analysis of the three tested *GBA* variants resulted in a significant P-value of 0.0050 (Figure 6, Supplementary Figure 14).

## Discussion

In the current study, we analysed multiple large datasets to examine whether genetic variants affect the penetrance and AAO of *GBA*-associated neurodegeneration. Our data show that at least 7% of the European PD population carries a common protein coding *GBA* risk variant versus ~4.5% of the general population. It is noteworthy that this number is an underestimation because in some datasets p.T369M and p.N370S were not genotyped or reliably imputed, and rarer disease-linked *GBA* variants were not captured. For LBD, these numbers are likely to be higher as seen in the NDRU-LBD dataset with ~13% of the LBD cases carrying a *GBA* variant.

One of the major findings of this study is that PD and LBD cases with a *GBA* variants often also carry a substantial number of other PD associated risk variants, which may modify disease risk and AAO. Pre-trial genetic analysis could assist in identifying those at higher risk for developing PD or LBD outside of the risk accounted for at the *GBA* target. Therefore, future clinical trials involving *GBA* carriers should include a comprehensive genetic assessment prior to enrolment, as we have previously suggested (Leonard et al. 2018). Additionally, disease predictions based on current genetic knowledge alone are not sufficient; larger datasets are needed and more non-genetic data, such as family history and smell test results can significantly improve disease prediction (Nalls et al. 2015c).

The link between PD and lysosomal pathways has already been established by both genetic and functional studies (Gan-Or et al. 2015b; Robak et al. 2017; Ysselstein et al. 2019). Moreover, in the latest PD GWAS (Nalls et al. 2018) numerous lysosomal and autophagy-related genes were associated with PD, emphasizing the central role of the lysosomal-autophagy pathway in disease. Interestingly, the two main loci identified that influence disease risk in *GBA* carriers are variants in close proximity to *SNCA* and *CTSB* (rs356219 and rs1293298), both implicated in the lysosomal autophagy pathway. The *SNCA* locus is well-studied and the current hypothesis is that variants in this locus increase *SNCA* expression of either total α-synuclein or specific mRNA isoforms, resulting in an increased risk of PD (Pihlstrøm et al. 2018; Soldner et al. 2016).

For the *CTSB* locus, no functional studies have been performed to dissect this locus in PD to date, although several genetic studies have nominated *CTSB* as causal gene in this region via either QTL analyses or Mendelian randomization (Li et al. 2019; Nalls et al. 2018). Therefore, it remains unclear whether *CTSB* is the actual causal gene in this locus. However, there is evidence linking *CTSB* to PD pathways. In this study, we see a decrease in processed Cathepsin B protein levels in iPSC-derived neurons from *GBA* variant carriers compared to non-carriers, suggesting a further reduction in lysosomal protease function in these cases. Depletion of *TMEM175*, another lysosomal gene associated with PD, significantly decreases protein expression and enzyme activity of Cathepsin B in rat hippocampal neurons (Sarah Jinn et al. 2017; S. Jinn et al. 2019). Cathepsin B has also been suggested to be involved in α-synuclein lysosomal degradation (McGlinchey and Lee 2015). Moreover, ceramide, which is one of the lipids produced by GCase, has been shown to be an activator of Cathepsin B-related pathways (F. Liu et al. 2016). Overall, these results fit with the general model that *GBA* risk variant carriers have impaired lysosomal function due to reduced or abolished GCase activity (Alcalay (Alcalay et al. 2015). The independent associations of the *SNCA* and *CTSB* loci with *GBA* associated PD suggest that increased *SNCA* expression and/or decreased *CTSB* mRNA expression further impair lysosomal function, resulting in an increased risk of PD. Additionally, we identified that Cathepsin B protein levels are reduced in p.N370S neurons compared to wildtype. Overall, these findings suggest that Cathepsin B might play a larger role in PD than previously thought and that increasing *CTSB/*Cathepsin B levels could be a potential therapeutic strategy. However, extensive functional *in vitro* and *in vivo* work is needed to validate these effects.

Despite the large datasets included in this study, it has several limitations. First, only one genome-wide significant locus was identified (*SNCA*), most likely due to the lack of power caused by the relatively low number of *GBA* variant carriers available. However, when we sought to reduce the burden of genome-wide correction by examining previously implicated PD risk loci, a significant association with the *CTSB* locus was identified. Second, the majority of the analyses were based on aggregated genotype data derived from several different genotyping arrays and centers/countries. As such, some datasets did not contain data on all three of the *GBA* variants of interest. It is also important to consider that different centers/countries might have used different measures of AAO or age at diagnosis and that included controls may develop PD at a later age. However, using the current data, we are still able to identify large effects, such as those driven by the GRS. Larger future initiatives specifically focused on *GBA* carriers will resolve these limitations and are needed to confirm these associations. Third, because the majority of the data included here were also used in the most recent PD GWAS meta-analysis (Nalls et al. 2018); there could be a bias towards the identification of known PD risk signals in this *GBA* dataset. We addressed this potential bias by looking for replication in several datasets that have not been included in the most recent PD GWAS meta-analysis, such as the UKB PD cases, the LBD dataset and the McGill *GBA* dataset. Unfortunately, we did not have a replication dataset for the *GBA* AAO results, so these results require replication in an independent dataset. Finally, some datasets might have excluded known *GBA* carriers prior to genotyping, which would result in an underestimation of total *GBA* carriers and the associated risk estimates. Of note, another relatively common *GBA* variant p.L444P (rs421016) is currently not imputable, since it is not included in the HRC reference panel and is in a complex region of the genome with a highly similar pseudogene.

Despite these limitations, several conclusions can be drawn from our results. First, the risk of PD in *GBA* carriers is influenced by specific variants at loci that are known to be associated more generally with PD risk, including *SNCA* and *CTSB*, the latter possibly through interaction with *GBA*. In addition, *GBA* associated risk is significantly affected by the GRS; however, the total contribution of the GRS is not large. In other words, genetic factors do not fully explain the partial penetrance of *GBA* variants in PD, hence environmental and aging-related factors probably contribute to *GBA* penetrance. Furthermore, variants in known PD loci, such as *SNCA* and *TMEM175*, as well as the GRS, might affect the AAO of *GBA*-associated parkinsonism at a similar scale compared to the effect seen in typical PD. Overall, these results highlight the importance of the lysosomal pathway in PD and nominates *CTSB* and *SNCA* as interesting candidate genes for functional analysis in both *GBA*-associated and general PD.

## Acknowledgements and Funding

We would like to thank all of the subjects who donated their time and biological samples to be part of this study. We would also like to thank all members of the International Parkinson Disease Genomics Consortium (IPDGC). For a complete overview of members, acknowledgements and funding, please see the Supplemental data and/or http://pdgenetics.org/partners. This work was supported in part by the Intramural Research Programs of the National Institute of Neurological Disorders and Stroke (NINDS), the National Institute on Aging (NIA), and the National Institute of Environmental Health Sciences both part of the National Institutes of Health, Department of Health and Human Services; project numbers 1ZIA-NS003154, Z01-AG000949-02 and Z01-ES101986. This work was also financially supported by the Michael J. Fox Foundation, the Canadian Consortium on Neurodegeneration in Aging (CCNA), the Canada First Research Excellence Fund (CFREF), awarded to McGill University for the Healthy Brains for Healthy Lives (HBHL) program. The Columbia University cohort is supported by the Parkinson’s Foundation, the National Institutes of Health [K02NS080915, and UL1 TR000040], and the Brookdale Foundation. The authors thank Ole Andreassen and the DemGene consortium for genotyping of the Oslo cohort. Samples used in the analyses presented in this article were obtained from the Golub Capital iPSC Parkinson’s Progression Markers Initiative (PPMI) Sub-study (www.ppmi-info.org/cell-lines). As such, the investigators within PPMI contributed to the design and implementation of PPMI and/or provided data and collected samples but did not participate in the analysis or writing of this report. For up-to-date information on the study, visit www.ppmi--info.org. Data used in the preparation of this article were obtained from the Parkinson’s Progression Markers Initiative (PPMI) database (www.ppmi-info.org/data). For up-to-date information on the study, visit www.ppmi-info.org. PPMI, a public-private partnership, is funded by the Michael J. Fox Foundation for Parkinson’s Research and funding partners, including AbbVie, Avid, Biogen, Bristol-Myers Squibb, Covance, GE Healthcare, Genentech, GlaxoSmithKline, Lilly, Lundbeck, Merck, Meso Scale Discovery, Pfizer, Piramal, Roche, Servier, Teva, UCB, and Golub Capital. Data and biospecimens used in preparation of this manuscript were obtained from the Parkinson’s Disease Biomarkers Program (PDBP) Consortium, part of the National Institute of Neurological Disorders and Stroke at the National Institutes of Health. Investigators include: Roger Albin, Roy Alcalay, Alberto Ascherio, DuBois Bowman, Alice Chen-Plotkin, Ted Dawson, Richard Dewey, Dwight German, Xuemei Huang, Rachel Saunders-Pullman, Liana Rosenthal, Clemens Scherzer, David Vaillancourt, Vladislav Petyuk, Andy West and Jing Zhang. The PDBP Investigators have not participated in reviewing the data analysis or content of the manuscript. DNA panels from the NINDS Human Genetics Resource Center DNA and Cell Line Repository (http://ccr.coriell.org/ninds) were used in this study, as well as clinical data. We thank the following brain banks for providing brain tissues: Banner Sun Health Research Institute, New York Brain Bank, Newcastle Brain Tissue Resource, University of Michigan Brain Bank, University of California San Diego Brain Bank, Duke University Brain Bank, Virginia Commonwealth University Brain Bank, and the Georgetown University Brain Bank. We would like to thank the NIA Baltimore Longitudinal Study of Aging for contributing tissue samples to the Johns Hopkins Alzheimer’s Disease Research Center. We are grateful to members of the North American Brain Expression Consortium (NABEC) for contributing DNA samples. This research has been conducted using the UK Biobank Resource under Application Number 33601. We would like to thank the NIH Neuro Brain Bank (https://neurobiobank.nih.gov) for contributing tissue samples. This study was supported in part by grants from the National Institutes of Health: U19-AG03365, P50 NS38377, and P50-AG005146. Tissue samples for genotyping were provided by the Johns Hopkins Morris K. Udall Center of Excellence for Parkinson’s Disease Research (NIH P50 NS38377) and the Johns Hopkins Alzheimer’s Disease Research Center. We are grateful for the support of the entire BIOCARD study team at Johns Hopkins University. Additionally, we acknowledge the contributions of the Geriatric Psychiatry Branch (GPB) in the intramural program of NIMH who initiated the BIOCARD study.

## Author contributions

Concept and design: Blauwendraat, Nalls, Gan-Or, Singleton

Statistical analysis: Blauwendraat, Nalls

Functional experiments: Reed

Contributed expertise, data or DNA samples: All

Drafting of the manuscript: Blauwendraat, Nalls, Gan-Or, Singleton

Critical revision of the manuscript for important intellectual content: All

## References

Abraham, Gad, Qiu, Yixuan, and Inouye, Michael (2017), ‘FlashPCA2: principal component analysis of Biobank-scale genotype datasets’, Bioinformatics, 33 (17), 2776–78.

Alcalay, Roy N., et al. (2015), ‘Glucocerebrosidase activity in Parkinson’s disease with and without GBA mutations’, Brain, 138 (Pt 9), 2648–58.

Anheim, M., et al. (2012), ‘Penetrance of Parkinson disease in glucocerebrosidase gene mutation carriers’, Neurology, 78 (6), 417–20.

Blauwendraat, C., et al. (2018a), ‘Frequency of Loss of Function Variants in LRRK2 in Parkinson Disease’, JAMA Neurol, 75 (11), 1416–22.

Blauwendraat, C., et al. (2019), ‘Parkinson’s disease age at onset genome-wide association study: Defining heritability, genetic loci, and alpha-synuclein mechanisms’, Mov Disord.

Blauwendraat, Cornelis, et al. (2018b), ‘Coding variation in GBA explains the majority of the SYT11-GBA Parkinson’s disease GWAS locus’, Mov. Disord., 33 (11), 1821–23.

Blauwendraat, Cornelis, et al. (2017), ‘NeuroChip, an updated version of the NeuroX genotyping platform to rapidly screen for variants associated with neurological diseases’, Neurobiol. Aging, 57, 247.e9–47.e13.

Butler, A., et al. (2018), ‘Integrating single-cell transcriptomic data across different conditions, technologies, and species’, Nat Biotechnol, 36 (5), 411–20.

Bycroft, Clare, et al. (2018), ‘The UK Biobank resource with deep phenotyping and genomic data’, Nature, 562 (7726), 203–09.

Consortium, G. TEx (2013), ‘The Genotype-Tissue Expression (GTEx) project’, Nat. Genet., 45 (6), 580–85.

Gan-Or, Z., et al. (2008), ‘Genotype-phenotype correlations between GBA mutations and Parkinson disease risk and onset’, Neurology, 70 (24), 2277–83.

Gan-Or, Z., et al. (2015a), ‘The Alzheimer disease BIN1 locus as a modifier of GBA-associated Parkinson disease’, J. Neurol., 262 (11), 2443–47.

Gan-Or, Ziv, Dion, Patrick A., and Rouleau, Guy A. (2015b), ‘Genetic perspective on the role of the autophagy-lysosome pathway in Parkinson disease’, Autophagy, 11 (9), 1443–57.

Gan-Or, Ziv, Liong, Christopher, and Alcalay, Roy N. (2018), ‘GBA-Associated Parkinson’s Disease and Other Synucleinopathies’, Curr. Neurol. Neurosci. Rep., 18 (8), 44.

Gan-Or, Ziv, et al. (2015c), ‘Differential effects of severe vs mild GBA mutations on Parkinson disease’, Neurology, 84 (9), 880–87.

Gibbs, J. R., et al. (2010), ‘Abundant quantitative trait loci exist for DNA methylation and gene expression in human brain’, PLoS Genet, 6 (5), e1000952.

Guerreiro, Rita, et al. (2018), ‘Investigating the genetic architecture of dementia with Lewy bodies: a two-stage genome-wide association study’, Lancet Neurol., 17 (1), 64–74.

Habib, N., et al. (2017), ‘Massively parallel single-nucleus RNA-seq with DroNc-seq’, Nat Methods, 14 (10), 955–58.

Jinn, S., et al. (2019), ‘Functionalization of the TMEM175 p.M393T Variant as a risk factor for Parkinson Disease’, Hum Mol Genet.

Jinn, Sarah, et al. (2017), ‘TMEM175 deficiency impairs lysosomal and mitochondrial function and increases α-synuclein aggregation’, Proc. Natl. Acad. Sci. U. S. A., 114 (9), 2389–94.

Leija-Salazar, Melissa, et al. (2019), ‘Evaluation of the detection of GBA missense mutations and other variants using the Oxford Nanopore MinION’, Mol Genet Genomic Med, e564.

Leonard, Hampton, et al. (2018), ‘Genetic variability and potential effects on clinical trial outcomes: perspectives in Parkinson’s disease’.

Lesage, Suzanne, et al. (2011), ‘Large-scale screening of the Gaucher’s disease-related glucocerebrosidase gene in Europeans with Parkinson’s disease’, Hum. Mol. Genet., 20 (1), 202–10.

Li, Y. I., et al. (2019), ‘Prioritizing Parkinson’s disease genes using population-scale transcriptomic data’, Nat Commun, 10 (1), 994.

Liu, B., et al. (2019), ‘Abundant associations with gene expression complicate GWAS follow-up’, Nat Genet, 51 (5), 768–69.

Liu, Feiyan, et al. (2016), ‘Ceramide activates lysosomal cathepsin B and cathepsin D to attenuate autophagy and induces ER stress to suppress myeloid-derived suppressor cells’, Oncotarget, 7 (51), 83907–25.

Malek, Naveed, et al. (2018), ‘Features of -associated Parkinson’s disease at presentation in the UK study’, J. Neurol. Neurosurg. Psychiatry, 89 (7), 702–09.

McGlinchey, Ryan P. and Lee, Jennifer C. (2015), ‘Cysteine cathepsins are essential in lysosomal degradation of α-synuclein’, Proc. Natl. Acad. Sci. U. S. A., 112 (30), 9322–27.

McKeith, I. G. (2006), ‘Consensus guidelines for the clinical and pathologic diagnosis of dementia with Lewy bodies (DLB): report of the Consortium on DLB International Workshop’, J. Alzheimers. Dis., 9 (3 Suppl), 417–23.

Nalls, Mike A., et al. (2015a), ‘Genetic risk and age in Parkinson’s disease: Continuum not stratum’, Mov. Disord., 30 (6), 850–54.

Nalls, Mike A., et al. (2015b), ‘NeuroX, a fast and efficient genotyping platform for investigation of neurodegenerative diseases’, Neurobiol. Aging, 36 (3), 1605.e7–12.

Nalls, Mike A., et al. (2015c), ‘Diagnosis of Parkinson’s disease on the basis of clinical and genetic classification: a population-based modelling study’, Lancet Neurol., 14 (10), 1002–09.

Nalls, Mike A., et al. (2018), ‘Parkinson’s disease genetics: identifying novel risk loci, providing causal insights and improving estimates of heritable risk’.

Pihlstrøm, Lasse, et al. (2018), ‘A comprehensive analysis of SNCA -related genetic risk in sporadic parkinson disease’, Ann. Neurol., 84 (1), 117–29.

Pruim, R. J., et al. (2010), ‘LocusZoom: regional visualization of genome-wide association scan results’, Bioinformatics, 26 (18), 2336–7.

Qi, Ting, et al. (2018), ‘Identifying gene targets for brain-related traits using transcriptomic and methylomic data from blood’, Nat. Commun., 9 (1), 2282.

Rana, Huma Q., et al. (2013), ‘Age-specific Parkinson disease risk in GBA mutation carriers: information for genetic counseling’, Genet. Med., 15 (2), 146–49.

Rivas, Manuel A., et al. (2018), ‘Insights into the genetic epidemiology of Crohn’s and rare diseases in the Ashkenazi Jewish population’, PLoS Genet., 14 (5), e1007329.

Robak, Laurie A., et al. (2017), ‘Excessive burden of lysosomal storage disorder gene variants in Parkinson’s disease’, Brain, 140 (12), 3191–203.

Robin, Xavier, et al. (2011), ‘pROC: an open-source package for R and S to analyze and compare ROC curves’, BMC Bioinformatics, 12 (1).

Ruskey, Jennifer A., et al. (2019), ‘Increased yield of full GBA sequencing in Ashkenazi Jews with Parkinson’s disease’, Eur. J. Med. Genet., 62 (1), 65–69.

Sidransky, E., et al. (2009), ‘Multicenter analysis of glucocerebrosidase mutations in Parkinson’s disease’, N. Engl. J. Med., 361 (17), 1651–61.

Soldner, Frank, et al. (2016), ‘Parkinson-associated risk variant in distal enhancer of α-synuclein modulates target gene expression’, Nature, 533 (7601), 95–99.

Sudlow, Cathie, et al. (2015), ‘UK biobank: an open access resource for identifying the causes of a wide range of complex diseases of middle and old age’, PLoS Med., 12 (3), e1001779.

Viechtbauer, W. (2010), ‘Conducting Meta-Analyses in R with the metafor Package’, Journal of Statistical Software, 36 (3), 1–48.

Wang, K., Li, M., and Hakonarson, H. (2010), ‘ANNOVAR: functional annotation of genetic variants from high-throughput sequencing data’, Nucleic Acids Res., 38 (16), e164–e64.

Willer, Cristen J., Li, Yun., and Abecasis, Gonçalo R. (2010), ‘METAL: fast and efficient meta-analysis of genomewide association scans’, Bioinformatics, 26 (17), 2190–91.

Yahalom, Gilad, et al. (2018), ‘Carriers of both GBA and LRRK2 mutations, compared to carriers of either, in Parkinson’s disease: Risk estimates and genotype-phenotype correlations’, Parkinsonism Relat. Disord.

Yang, J., et al. (2012), ‘Conditional and joint multiple-SNP analysis of GWAS summary statistics identifies additional variants influencing complex traits’, Nat Genet, 44 (4), 369–75, S1-3.

Yang, Jian, et al. (2011), ‘GCTA: a tool for genome-wide complex trait analysis’, Am. J. Hum. Genet., 88 (1), 76–82.

Ysselstein, Daniel, Shulman, Joshua M., and Krainc, Dimitri (2019), ‘Emerging links between pediatric lysosomal storage diseases and adult parkinsonism’, Mov. Disord.

Zhan, Xiaowei, et al. (2016), ‘RVTESTS: an efficient and comprehensive tool for rare variant association analysis using sequence data’, Bioinformatics, 32 (9), 1423–26.

