## Supplement for "Genetic modifiers of risk and age at onset in *GBA* associated Parkinson’s disease and Lewy body dementia": GBA modifiers Supplemental data final version.docx

Blauwendraat et al. 2019

***Supplemental Data***

**Detail description of processing of genotyping data**

**Sequencing validation of GBA variants**

**Processing of genotyping data**

Genotyping data (all Illumina platform based) was obtained from International Parkinson’s Disease Genomics Consortium (IPDGC) members, collaborators and public resources (Supplementary Table 2 and 3). All datasets underwent quality control separately, both on individual level data and variant level data before imputation as previously described (Blauwendraat et al. 2018; Nalls et al. 2018). In brief, we excluded individuals with discordance between genetic and reported sex, low call rate (<95%), heterozygosity outliers (F statistic cut-off of > -0.15 and < 0.15), ancestry outliers (+/- 6 standard deviations from means of eigenvectors 1 and 2 of the 1000 Genomes phase 3 CEU and TSI populations from principal components(Genomes Project et al. 2015), and individuals with cryptic relatedness of > 0.125 using PLINK (version 1.9) (Chang et al. 2015) or GCTA (version 1.02) (Yang et al. 2011) keeping a random proband from any related groups. We further excluded genotypes with a missingness rate of > 5%, a minor allele frequency < 0.001 and Hardy-Weinberg equilibrium of P =>1E-6. Remaining samples were submitted per dataset separately to the Michigan Imputation Server (Das et al. 2016) using Eagle v2.3 imputation (Loh et al. 2016) based on reference panel HRC r1.1 2016 (McCarthy et al. 2016). After imputation, all datasets were merged and analysed for duplicates. One individual was removed at random from pairs of duplicated or related individuals. Age at onset was defined based on patient report of initial manifestation of parkinsonian motor signs (tremor, bradykinesia, rigidity or gait impairment). When this information was not available, age of diagnosis was used as a proxy for onset age, which previously has been shown to be highly correlated with age of onset (Blauwendraat et al. 2018).

Principal components (PCs) were created on directly assayed genotypes overlapping in all datasets using PLINK. For the PC calculation, variants were filtered for minor allele frequency (>0.05), genotype missingness (<0.25) and Hardy–Weinberg equilibrium (P =>1E-6). Remaining variants were pruned using a PLINK pairwise pruning (default settings, window size of 50, window shift: 5 SNPs and r2 of 0.5) and the additional following high LD regions were excluded chr5:44000000-51500000, chr6:25000000-33500000, chr8:8000000-12000000 and chr11:45000000-57000000. PCs were calculated on the pruned variants.

**Sequencing validation of *GBA* variants**

In order to validate the genotyping and imputation of *GBA* variants we mined sequencing data from two parallel IPDGC projects. In total, 144 *GBA* carriers (17 controls and 127 cases) were genome sequenced and 184 *GBA* carriers (55 controls and 129 cases) were targeted resequenced, with 9 *GBA* carriers overlapping between both, bringing the total to 319 *GBA* carriers (65 controls and 254 cases). All 49 p.N369S carriers were confirmed by sequencing of which 47 were genotyped and 2 were imputed. One heterozygous carrier in genotyped data appeared to be a homozygous carrier in the genome sequence data. In total 79 p.T369M heterozygous carriers were confirmed by sequencing and all were directly genotyped. 190 out of 193 p.E326K carriers were confirmed by sequencing of which 63 were directly genotyped and 130 were imputed. Four p.E326K carriers were homozygous in the sequence data of which two were confirmed and two were not (all imputed). Three p.E326K carriers were imputed to be heterozygous but could not be confirmed in the sequence data. Additionally, the majority of the PROBAND cohort was Sanger sequenced. Of the 89 p.E326K imputed carriers 86 were Sanger sequenced and 81 were confirmed (94%). Additionally, both p.N369S and p.T369M were not properly imputed (R^2^<0.8) in the PROBAND and therefore by Sanger sequencing we identified 10 additional p.N369S carriers and 33 p.T369M carriers. Overall, the concordance rate between the imputed/genotype data and the sequence data for the presence of at least one p.N369S and p.T369M allele was 100% and for p.E326K was 98% and 94% in the next-generation and Sanger sequencing data, respectively. *GBA* carriers were corrected or excluded if they could not be validated from the sequencing data in subsequent analysis.

**Whole genome sequencing**

Whole genome sequencing (WGS) data was generated at the Laboratory of Neurogenetics (LNG) at the National Institutes of Health from multiple cohorts. Paired-end read sequences were processed in accordance with the standard pipeline developed by the Centers for Common Disease Genomics (https://www.genome.gov/27563570/). This standard allows for whole-genome sequence (WGS) processed by different groups or centers to generate "functionally equivalent" (FE) results (Regier et al. 2018). The GRCh38DH reference genome was used for alignment as specified in the FE standard (https://github.com/CCDG/Pipeline-Standardization/blob/master/PipelineStandard.md). The Broad Institute’s implementation of this FE standardized pipeline, which incorporates the GATK (2016) Best Practices (Van der Auwera et al. 2013), is publicly available (https://github.com/gatk-workflows/broad-prod-wgs-germline-snps-indels) and was used for WGS processing. Single-nucleotide (SNV) and InDel variants were called from the processed WGS data following the GATK (2016) Best Practices using the Broad Institute’s workflow for joint discovery and Variant Quality Score Recalibration (VQSR) (https://github.com/CCDG/Pipeline-Standardization/blob/master/PipelineStandard.md).

For quality control each sample is checked using common methods for genotypes as well and sequence related metrics. Using PLINK v1.9 (Chang et al. 2015) each sample’s genotype missingness rate (included in this study if < 95%), heterozygosity rate (exceeding +/- 0.15 F-stat was an exclusion criteria here), sex concordance between genetically ascertained and clinically scertained sex status were used in initial sample filtering and excessive relatedness (>0.125) and having non-European ancestry based on a principal components analysis with the Hapmap Genome reference panel (International HapMap et al. 2010). The King v2.1.3 kinship tool (Manichaikul et al. 2010) was used to check for the presence of duplicate samples and to check concordance with existing genotypes for subjects for whom data are available. Sequence and alignment related metrics generated by the Broad’s implementation of the FE standardized pipeline were inspected for potential quality problems. This included the sample’s mean sequence depth (< 30X as an exclusion criteria) and contamination rate (> 2%, also an exclusion criteria), as reported by VerifyBamID (Jun et al. 2012), and SNV count as reported by Picard’s CollectVariantCallingMetrics (< 3 StDev resulted in sample exclusion) based on the sample’s genomic vcf (gvcf). Subsequent genotypes were filtered to exclude minor allele frequency of less than 0.01, missing rate more than 5%, and HWE threshold of 1.0E-4 in controls.

**References**

Blauwendraat, Cornelis, et al. (2018), 'Parkinson disease age of onset GWAS: defining heritability, genetic loci and a-synuclein mechanisms'.

Chang, Christopher C., et al. (2015), 'Second-generation PLINK: rising to the challenge of larger and richer datasets', *Gigascience,* 4, 7.

Das, Sayantan, et al. (2016), 'Next-generation genotype imputation service and methods', *Nat. Genet.,* 48 (10), 1284-87.

Genomes Project, Consortium, et al. (2015), 'A global reference for human genetic variation', *Nature,* 526 (7571), 68-74.

International HapMap, Consortium, et al. (2010), 'Integrating common and rare genetic variation in diverse human populations', *Nature,* 467 (7311), 52-8.

Jun, G., et al. (2012), 'Detecting and estimating contamination of human DNA samples in sequencing and array-based genotype data', *Am J Hum Genet,* 91 (5), 839-48.

Loh, Po-Ru, et al. (2016), 'Reference-based phasing using the Haplotype Reference Consortium panel', *Nat. Genet.,* 48 (11), 1443-48.

Manichaikul, A., et al. (2010), 'Robust relationship inference in genome-wide association studies', *Bioinformatics,* 26 (22), 2867-73.

McCarthy, Shane, et al. (2016), 'A reference panel of 64,976 haplotypes for genotype imputation', *Nat. Genet.,* 48 (10), 1279-83.

Nalls, Mike A., et al. (2018), 'Parkinson's disease genetics: identifying novel risk loci, providing causal insights and improving estimates of heritable risk'.

Regier, A. A., et al. (2018), 'Functional equivalence of genome sequencing analysis pipelines enables harmonized variant calling across human genetics projects', *Nat Commun,* 9 (1), 4038.

Van der Auwera, G. A., et al. (2013), 'From FastQ data to high confidence variant calls: the Genome Analysis Toolkit best practices pipeline', *Curr Protoc Bioinformatics,* 43, 11 10 1-33.

Yang, Jian, et al. (2011), 'GCTA: a tool for genome-wide complex trait analysis', *Am. J. Hum. Genet.,* 88 (1), 76-82.

**IPDGC consortium members and affiliations:**

**United Kingdom:** Alastair J Noyce (Preventive Neurology Unit, Wolfson Institute of Preventive Medicine, QMUL, London, UK and Department of Molecular Neuroscience, UCL, London, UK), Rauan Kaiyrzhanov (Department of Molecular Neuroscience, UCL Institute of Neurology, London, UK), Ben Middlehurst (Institute of Translational Medicine, University of Liverpool, Liverpool, UK), Demis A Kia (UCL Genetics Institute; and Department of Molecular Neuroscience, UCL Institute of Neurology, London, UK), Manuela Tan (Department of Clinical Neuroscience, University College London, London, UK), Henry Houlden (Department of Molecular Neuroscience, UCL Institute of Neurology, London, UK), Huw R Morris (Department of Clinical Neuroscience, University College London, London, UK), Helene Plun-Favreau (Department of Molecular Neuroscience, UCL Institute of Neurology, London, UK), Peter Holmans (Biostatistics & Bioinformatics Unit, Institute of Psychological Medicine and Clinical Neuroscience, MRC Centre for Neuropsychiatric Genetics & Genomics, Cardiff, UK), John Hardy (Department of Molecular Neuroscience, UCL Institute of Neurology, London, UK), Daniah Trabzuni (Department of Molecular Neuroscience, UCL Institute of Neurology, London, UK; Department of Genetics, King Faisal Specialist Hospital and Research Centre, Riyadh, 11211 Saudi Arabia), John Quinn PhD (Institute of Translational Medicine, University of Liverpool, Liverpool, UK), Kin Y Mok (Department of Molecular Neuroscience, UCL Institute of Neurology, London, UK), Kerri J. Kinghorn (Institute of Healthy Ageing, University College London, London, UK), Kimberley Billingsley (Institute of Translational Medicine, University of Liverpool, Liverpool, UK), Nicholas W Wood (UCL Genetics Institute; and Department of Molecular Neuroscience, UCL Institute of Neurology, London, UK), Patrick Lewis (University of Reading, Reading, UK), Sebastian Schreglmann (Department of Molecular Neuroscience, UCL Institute of Neurology, London, UK), Ruth Lovering (University College London, London, UK), Lea R’Bibo (Department of Molecular Neuroscience, UCL Institute of Neurology, London, UK), Claudia Manzoni (University of Reading, Reading, UK), Mie Rizig (Department of Molecular Neuroscience, UCL Institute of Neurology, London, UK), Mina Ryten (Department of Molecular Neuroscience, UCL Institute of Neurology, London, UK), Sebastian Guelfi (Department of Molecular Neuroscience, UCL Institute of Neurology, London, UK), Valentina Escott-Price (MRC Centre for Neuropsychiatric Genetics and Genomics, Cardiff University School of Medicine, Cardiff, UK), Viorica Chelban (Department of Molecular Neuroscience, UCL Institute of Neurology, London, UK), Thomas Foltynie (UCL Institute of Neurology, London, UK), Nigel Williams (MRC Centre for Neuropsychiatric Genetics and Genomics, Cardiff, UK), Karen E. Morrison (Faculty of Medicine, University of Southampton, UK), Carl Clarke (University of Birmingham, Birmingham, UK and Sandwell and West Birmingham Hospitals NHS Trust, Birmingham, UK).

**France:** Alexis Brice (Institut du Cerveau et de la Moelle épinière, ICM, Inserm U 1127, CNRS, UMR 7225, Sorbonne Universités, UPMC University Paris 06, UMR S 1127, AP-HP, Pitié-Salpêtrière Hospital, Paris, France), Fabrice Danjou (Institut du Cerveau et de la Moelle épinière, ICM, Inserm U 1127, CNRS, UMR 7225, Sorbonne Universités, UPMC University Paris 06, UMR S 1127, AP-HP, Pitié-Salpêtrière Hospital, Paris, France), Suzanne Lesage (Institut du Cerveau et de la Moelle épinière, ICM, Inserm U 1127, CNRS, UMR 7225, Sorbonne Universités, UPMC University Paris 06, UMR S 1127, AP-HP, Pitié-Salpêtrière Hospital, Paris, France), Jean-Christophe Corvol (Institut du Cerveau et de la Moelle épinière, ICM, Inserm U 1127, CNRS, UMR 7225, Sorbonne Universités, UPMC University Paris 06, UMR S 1127, Centre d’Investigation Clinique Pitié Neurosciences CIC-1422, AP-HP, Pitié-Salpêtrière Hospital, Paris, France), Maria Martinez (INSERM UMR 1220; and Paul Sabatier University, Toulouse, France),

**Germany:** Claudia Schulte (Department for Neurodegenerative Diseases, Hertie Institute for Clinical Brain Research, University of Tübingen, and DZNE, German Center for Neurodegenerative Diseases, Tübingen, Germany), Kathrin Brockmann (Department for Neurodegenerative Diseases, Hertie Institute for Clinical Brain Research, University of Tübingen, and DZNE, German Center for Neurodegenerative Diseases, Tübingen, Germany), Javier Simón-Sánchez (Department for Neurodegenerative Diseases, Hertie Institute for Clinical Brain Research, University of Tübingen, and DZNE, German Center for Neurodegenerative Diseases, Tübingen, Germany), Peter Heutink (DZNE, German Center for Neurodegenerative Diseases and Department for Neurodegenerative Diseases, Hertie Institute for Clinical Brain Research, University of Tübingen, Tübingen, Germany), Patrizia Rizzu (DZNE, German Center for Neurodegenerative Diseases), Manu Sharma (Centre for Genetic Epidemiology, Institute for Clinical Epidemiology and Applied Biometry, University of Tubingen, Germany), Thomas Gasser (Department for Neurodegenerative Diseases, Hertie Institute for Clinical Brain Research, and DZNE, German Center for Neurodegenerative Diseases, Tübingen, Germany),

**United States of America:** Mark R Cookson (Laboratory of Neurogenetics, National Institute on Aging, Bethesda, USA), Sara Bandres-Ciga (Laboratory of Neurogenetics, National Institute on Aging, Bethesda, MD, USA), Cornelis Blauwendraat (National Institute on Aging and National Institute of Neurological Disorders and Stroke, USA), David W. Craig (Department of Translational Genomics, Keck School of Medicine, University of Southern California, Los Angeles, USA), Derek Narendra (Inherited Movement Disorders Unit, National Institute of Neurological Disorders and Stroke, Bethesda, MD, USA), Faraz Faghri (Laboratory of Neurogenetics, National Institute on Aging, Bethesda, USA; Department of Computer Science, University of Illinois at Urbana-Champaign, Urbana, IL, USA), J Raphael Gibbs (Laboratory of Neurogenetics, National Institute on Aging, National Institutes of Health, Bethesda, MD, USA), Dena G. Hernandez (Laboratory of Neurogenetics, National Institute on Aging, Bethesda, MD, USA), Kendall Van Keuren-Jensen (Neurogenomics Division, TGen, Phoenix, AZ USA), Joshua M. Shulman (Departments of Neurology, Neuroscience, and Molecular & Human Genetics, Baylor College of Medicine, Houston, Texas, USA; Jan and Dan Duncan Neurological Research Institute, Texas Children’s Hospital, Houston, Texas, USA), Hirotaka Iwaki (Laboratory of Neurogenetics, National Institute on Aging, Bethesda, MD, USA), Hampton L. Leonard (Laboratory of Neurogenetics, National Institute on Aging, Bethesda, MD, USA), Mike A. Nalls (Laboratory of Neurogenetics, National Institute on Aging, Bethesda, USA; CEO/Consultant Data Tecnica International, Glen Echo, MD, USA), Laurie Robak (Baylor College of Medicine, Houston, Texas, USA), Jose Bras (Center for Neurodegenerative Science, Van Andel Research Institute, Grand Rapids, Michigan, USA), Rita Guerreiro (Center for Neurodegenerative Science, Van Andel Research Institute, Grand Rapids, Michigan, USA), Steven Lubbe (Ken and Ruth Davee Department of Neurology, Northwestern University Feinberg School of Medicine, Chicago, IL, USA), Steven Finkbeiner (Departments of Neurology and Physiology, University of California, San Francisco; Gladstone Institute of Neurological Disease; Taube/Koret Center for Neurodegenerative Disease Research, San Francisco, CA, USA), Niccolo E. Mencacci (Northwestern University Feinberg School of Medicine, Chicago, IL, USA), Codrin Lungu (National Institutes of Health Division of Clinical Research, NINDS, National Institutes of Health, Bethesda, MD, USA), Andrew B Singleton (Laboratory of Neurogenetics, National Institute on Aging, Bethesda, MD, USA), Sonja W. Scholz (Neurodegenerative Diseases Research Unit, National Institute of Neurological Disorders and Stroke, Bethesda, MD, USA), Xylena Reed (Laboratory of Neurogenetics, National Institute on Aging, Bethesda, MD, USA). Roy N. Alcalay (Department of Neurology, College of Physicians and Surgeons, Columbia University Medical Center, New York, NY, USA, Taub Institute for Research on Alzheimer's Disease and the Aging Brain, College of Physicians and Surgeons, Columbia University Medical Center, New York, NY, USA)

**Canada:** Ziv Gan-Or (Montreal Neurological Institute and Hospital, Department of Neurology & Neurosurgery, Department of Human Genetics, McGill University, Montréal, QC, H3A 0G4, Canada), Guy A. Rouleau (Montreal Neurological Institute and Hospital, Department of Neurology & Neurosurgery, Department of Human Genetics, McGill University, Montréal, QC, H3A 0G4, Canada), Lynne Krohn (Montreal Neurological Institute and Hospital, Department of Neurology & Neurosurgery, Department of Human Genetics, McGill University, Montréal, QC, H3A 0G4, Canada), Lynne Krohn (Montreal Neurological Institute and Hospital, Department of Neurology & Neurosurgery, Department of Human Genetics, McGill University, Montréal, QC, H3A 0G4, Canada),

**The Netherlands:** Jacobus J van Hilten (Department of Neurology, Leiden University Medical Center, Leiden, Netherlands), Johan Marinus (Department of Neurology, Leiden University Medical Center, Leiden, Netherlands)

**Spain:** Astrid D. Adarmes-Gómez (Instituto de Biomedicina de Sevilla (IBiS), Hospital Universitario Virgen del Rocío/CSIC/Universidad de Sevilla, Seville), Miquel Aguilar (Fundació Docència i Recerca Mútua de Terrassa and Movement Disorders Unit, Department of Neurology, University Hospital Mutua de Terrassa, Terrassa, Barcelona.), Ignacio Alvarez (Fundació Docència i Recerca Mútua de Terrassa and Movement Disorders Unit, Department of Neurology, University Hospital Mutua de Terrassa, Terrassa, Barcelona.),Victoria Alvarez (Hospital Universitario Central de Asturias, Oviedo), Francisco Javier Barrero (Hospital Universitario San Cecilio de Granada, Universidad de Granada), Jesús Alberto Bergareche Yarza (Instituto de Investigación Sanitaria Biodonostia, San Sebastián), Inmaculada Bernal-Bernal (Instituto de Biomedicina de Sevilla (IBiS), Hospital Universitario Virgen del Rocío/CSIC/Universidad de Sevilla, Seville), Marta Blazquez (Hospital Universitario Central de Asturias, Oviedo), Marta Bonilla-Toribio (Instituto de Biomedicina de Sevilla (IBiS), Hospital Universitario Virgen del Rocío/CSIC/Universidad de Sevilla, Seville), Juan A. Botía (Universidad de Murcia, Murcia), María Teresa Boungiorno (Fundació Docència i Recerca Mútua de Terrassa and Movement Disorders Unit, Department of Neurology, University Hospital Mutua de Terrassa, Terrassa, Barcelona.) Dolores Buiza-Rueda (Instituto de Biomedicina de Sevilla (IBiS), Hospital Universitario Virgen del Rocío/CSIC/Universidad de Sevilla, Seville), Ana Cámara (Hospital Clinic de Barcelona), Fátima Carrillo (Instituto de Biomedicina de Sevilla (IBiS), Hospital Universitario Virgen del Rocío/CSIC/Universidad de Sevilla, Seville), Mario Carrión-Claro (Instituto de Biomedicina de Sevilla (IBiS), Hospital Universitario Virgen del Rocío/CSIC/Universidad de Sevilla, Seville), Debora Cerdan (Hospital General de Segovia, Segovia), Jordi Clarimón (Memory Unit, Department of Neurology, IIB Sant Pau, Hospital de la Santa Creu i Sant Pau, Universitat Autònoma de Barcelona and Centro de Investigación Biomédica en Red en Enfermedades Neurodegenerativas (CIBERNED), Madrid),Yaroslau Compta (Hospital Clinic de Barcelona), Monica Diez-Fairen (Fundació Docència i Recerca Mútua de Terrassa and Movement Disorders Unit, Department of Neurology, University Hospital Mutua de Terrassa, Terrassa, Barcelona.), Oriol Dols-Icardo (Memory Unit, Department of Neurology, IIB Sant Pau, Hospital de la Santa Creu i Sant Pau, Universitat Autònoma de Barcelona, Barcelona, and Centro de Investigación Biomédica en Red en Enfermedades Neurodegenerativas (CIBERNED), Madrid), Jacinto Duarte (Hospital General de Segovia, Segovia), Raquel Duran (Centro de Investigacion Biomedica, Universidad de Granada, Granada), Francisco Escamilla-Sevilla (Hospital Universitario Virgen de las Nieves, Instituto de Investigación Biosanitaria de Granada, Granada), Mario Ezquerra (Hospital Clinic de Barcelona), Cici Feliz (Departmento de Neurologia, Instituto de Investigación Sanitaria Fundación Jiménez Díaz, Madrid, Spain), Manel Fernández (Hospital Clinic de Barcelona), Rubén Fernández-Santiago (Hospital Clinic de Barcelona), Ciara Garcia (Hospital Universitario Central de Asturias, Oviedo), Pedro García-Ruiz (Instituto de Investigación Sanitaria Fundación Jiménez Díaz, Madrid), Pilar Gómez-Garre (Instituto de Biomedicina de Sevilla (IBiS), Hospital Universitario Virgen del Rocío/CSIC/Universidad de Sevilla, Seville), Maria Jose Gomez Heredia (Hospital Universitario Virgen de la Victoria, Malaga), Isabel Gonzalez-Aramburu (Hospital Universitario Marqués de Valdecilla-IDIVAL, Santander) ,Ana Gorostidi Pagola (Instituto de Investigación Sanitaria Biodonostia, San Sebastián), Janet Hoenicka (Institut de Recerca Sant Joan de Déu, Barcelona), Jon Infante (Hospital Universitario Marqués de Valdecilla-IDIVAL and University of Cantabria, Santander, and Centro de Investigación Biomédica en Red en Enfermedades Neurodegenerativas (CIBERNED), Silvia Jesús (Instituto de Biomedicina de Sevilla (IBiS), Hospital Universitario Virgen del Rocío/CSIC/Universidad de Sevilla, Seville), Adriano Jimenez-Escrig (Hospital Universitario Ramón y Cajal, Madrid), Jaime Kulisevsky (Movement Disorders Unit, Department of Neurology, IIB Sant Pau, Hospital de la Santa Creu i Sant Pau, Universitat Autònoma de Barcelona, Barcelona, and Centro de Investigación Biomédica en Red en Enfermedades Neurodegenerativas (CIBERNED)), Miguel A. Labrador-Espinosa (Instituto de Biomedicina de Sevilla (IBiS), Hospital Universitario Virgen del Rocío/CSIC/Universidad de Sevilla, Seville), Jose Luis Lopez-Sendon (Hospital Universitario Ramón y Cajal, Madrid), Adolfo López de Munain Arregui (Instituto de Investigación Sanitaria Biodonostia, San Sebastián), Daniel Macias (Instituto de Biomedicina de Sevilla (IBiS), Hospital Universitario Virgen del Rocío/CSIC/Universidad de Sevilla, Seville), Irene Martínez Torres (Department of Neurology, Instituto de Investigación Sanitaria La Fe, Hospital Universitario y Politécnico La Fe, Valencia), Juan Marín (Movement Disorders Unit, Department of Neurology, IIB Sant Pau, Hospital de la Santa Creu i Sant Pau, Universitat Autònoma de Barcelona, Barcelona, and Centro de Investigación Biomédica en Red en Enfermedades Neurodegenerativas (CIBERNED)), Maria Jose Marti (Hospital Clinic Barcelona), Juan Carlos Martínez-Castrillo (Instituto Ramón y Cajal de Investigación Sanitaria, Hospital Universitario Ramón y Cajal, Madrid), Carlota Méndez-del-Barrio (Instituto de Biomedicina de Sevilla (IBiS), Hospital Universitario Virgen del Rocío/CSIC/Universidad de Sevilla, Seville), Manuel Menéndez González (Hospital Universitario Central de Asturias, Oviedo), Marina Mata (Department of Neurology, Hospital Universitario Infanta Sofía, Madrid, Spain) Adolfo Mínguez (Hospital Universitario Virgen de las Nieves, Granada, Instituto de Investigación Biosanitaria de Granada), Pablo Mir (Instituto de Biomedicina de Sevilla (IBiS), Hospital Universitario Virgen del Rocío/CSIC/Universidad de Sevilla, Seville), Elisabet Mondragon Rezola (Instituto de Investigación Sanitaria Biodonostia, San Sebastián), Esteban Muñoz (Hospital Clinic Barcelona), Javier Pagonabarraga (Movement Disorders Unit, Department of Neurology, IIB Sant Pau, Hospital de la Santa Creu i Sant Pau, Universitat Autònoma de Barcelona, Barcelona, and Centro de Investigación Biomédica en Red en Enfermedades Neurodegenerativas (CIBERNED)), Pau Pastor (Fundació Docència i Recerca Mútua de Terrassa and Movement Disorders Unit, Department of Neurology, University Hospital Mutua de Terrassa, Terrassa, Barcelona.), Francisco Perez Errazquin (Hospital Universitario Virgen de la Victoria, Malaga), Teresa Periñán-Tocino (Instituto de Biomedicina de Sevilla (IBiS), Hospital Universitario Virgen del Rocío/CSIC/Universidad de Sevilla, Seville), Javier Ruiz-Martínez (Hospital Universitario Donostia, Instituto de Investigación Sanitaria Biodonostia, San Sebastián), Clara Ruz (Centro de Investigacion Biomedica, Universidad de Granada, Granada), Antonio Sanchez Rodriguez (Hospital Universitario Marqués de Valdecilla-IDIVAL, Santander), María Sierra (Hospital Universitario Marqués de Valdecilla-IDIVAL, Santander), Esther Suarez-Sanmartin (Hospital Universitario Central de Asturias, Oviedo), Cesar Tabernero (Hospital General de Segovia, Segovia), Juan Pablo Tartari (Fundació Docència i Recerca Mútua de Terrassa and Movement Disorders Unit, Department of Neurology, University Hospital Mutua de Terrassa, Terrassa, Barcelona), Cristina Tejera-Parrado (Instituto de Biomedicina de Sevilla (IBiS), Hospital Universitario Virgen del Rocío/CSIC/Universidad de Sevilla, Seville), Eduard Tolosa (Hospital Clinic Barcelona), Francesc Valldeoriola (Hospital Clinic Barcelona), Laura Vargas-González (Instituto de Biomedicina de Sevilla (IBiS), Hospital Universitario Virgen del Rocío/CSIC/Universidad de Sevilla, Seville), Lydia Vela (Department of Neurology, Hospital Universitario Fundación Alcorcón, Madrid), Francisco Vives (Centro de Investigacion Biomedica, Universidad de Granada, Granada).

**Austria:** Alexander Zimprich (Department of Neurology, Medical University of Vienna, Austria)

**Norway:** Lasse Pihlstrom (Department of Neurology, Oslo University Hospital, Oslo, Norway), Mathias Toft (Department of Neurology and Institute of Clinical Medicine, Oslo University Hospital, Oslo, Norway)

**Estonia**: Sulev Koks (Department of Pathophysiology, University of Tartu, Tartu, Estonia; Department of Reproductive Biology, Estonian University of Life Sciences, Tartu, Estonia), Pille Taba (Department of Neurology and Neurosurgery, University of Tartu, Tartu, Estonia)

**Israel:** Sharon Hassin-Baer (The Movement Disorders Institute, Department of Neurology and Sagol Neuroscience Center, Chaim Sheba Medical Center, Tel-Hashomer, 5262101, Ramat Gan, Israel, Sackler Faculty of Medicine, Tel Aviv University, Tel Aviv, Israel)**IPDGC consortium members and affiliations:**

**United Kingdom:** Alastair J Noyce (Preventive Neurology Unit, Wolfson Institute of Preventive Medicine, QMUL, London, UK and Department of Molecular Neuroscience, UCL, London, UK), Rauan Kaiyrzhanov (Department of Molecular Neuroscience, UCL Institute of Neurology, London, UK), Ben Middlehurst (Institute of Translational Medicine, University of Liverpool, Liverpool, UK), Demis A Kia (UCL Genetics Institute; and Department of Molecular Neuroscience, UCL Institute of Neurology, London, UK), Manuela Tan (Department of Clinical Neuroscience, University College London, London, UK), Henry Houlden (Department of Molecular Neuroscience, UCL Institute of Neurology, London, UK), Huw R Morris (Department of Clinical Neuroscience, University College London, London, UK), Helene Plun-Favreau (Department of Molecular Neuroscience, UCL Institute of Neurology, London, UK), Peter Holmans (Biostatistics & Bioinformatics Unit, Institute of Psychological Medicine and Clinical Neuroscience, MRC Centre for Neuropsychiatric Genetics & Genomics, Cardiff, UK), John Hardy (Department of Molecular Neuroscience, UCL Institute of Neurology, London, UK), Daniah Trabzuni (Department of Molecular Neuroscience, UCL Institute of Neurology, London, UK; Department of Genetics, King Faisal Specialist Hospital and Research Centre, Riyadh, 11211 Saudi Arabia), John Quinn PhD (Institute of Translational Medicine, University of Liverpool, Liverpool, UK), Kin Y Mok (Department of Molecular Neuroscience, UCL Institute of Neurology, London, UK), Kerri J. Kinghorn (Institute of Healthy Ageing, University College London, London, UK), Kimberley Billingsley (Institute of Translational Medicine, University of Liverpool, Liverpool, UK), Nicholas W Wood (UCL Genetics Institute; and Department of Molecular Neuroscience, UCL Institute of Neurology, London, UK), Patrick Lewis (University of Reading, Reading, UK), Sebastian Schreglmann (Department of Molecular Neuroscience, UCL Institute of Neurology, London, UK), Ruth Lovering (University College London, London, UK), Lea R’Bibo (Department of Molecular Neuroscience, UCL Institute of Neurology, London, UK), Claudia Manzoni (University of Reading, Reading, UK), Mie Rizig (Department of Molecular Neuroscience, UCL Institute of Neurology, London, UK), Mina Ryten (Department of Molecular Neuroscience, UCL Institute of Neurology, London, UK), Sebastian Guelfi (Department of Molecular Neuroscience, UCL Institute of Neurology, London, UK), Valentina Escott-Price (MRC Centre for Neuropsychiatric Genetics and Genomics, Cardiff University School of Medicine, Cardiff, UK), Viorica Chelban (Department of Molecular Neuroscience, UCL Institute of Neurology, London, UK), Thomas Foltynie (UCL Institute of Neurology, London, UK), Nigel Williams (MRC Centre for Neuropsychiatric Genetics and Genomics, Cardiff, UK), Karen E. Morrison (Faculty of Medicine, University of Southampton, UK), Carl Clarke (University of Birmingham, Birmingham, UK and Sandwell and West Birmingham Hospitals NHS Trust, Birmingham, UK).

**France:** Alexis Brice (Institut du Cerveau et de la Moelle épinière, ICM, Inserm U 1127, CNRS, UMR 7225, Sorbonne Universités, UPMC University Paris 06, UMR S 1127, AP-HP, Pitié-Salpêtrière Hospital, Paris, France), Fabrice Danjou (Institut du Cerveau et de la Moelle épinière, ICM, Inserm U 1127, CNRS, UMR 7225, Sorbonne Universités, UPMC University Paris 06, UMR S 1127, AP-HP, Pitié-Salpêtrière Hospital, Paris, France), Suzanne Lesage (Institut du Cerveau et de la Moelle épinière, ICM, Inserm U 1127, CNRS, UMR 7225, Sorbonne Universités, UPMC University Paris 06, UMR S 1127, AP-HP, Pitié-Salpêtrière Hospital, Paris, France), Jean-Christophe Corvol (Institut du Cerveau et de la Moelle épinière, ICM, Inserm U 1127, CNRS, UMR 7225, Sorbonne Universités, UPMC University Paris 06, UMR S 1127, Centre d’Investigation Clinique Pitié Neurosciences CIC-1422, AP-HP, Pitié-Salpêtrière Hospital, Paris, France), Maria Martinez (INSERM UMR 1220; and Paul Sabatier University, Toulouse, France),

**Germany:** Claudia Schulte (Department for Neurodegenerative Diseases, Hertie Institute for Clinical Brain Research, University of Tübingen, and DZNE, German Center for Neurodegenerative Diseases, Tübingen, Germany), Kathrin Brockmann (Department for Neurodegenerative Diseases, Hertie Institute for Clinical Brain Research, University of Tübingen, and DZNE, German Center for Neurodegenerative Diseases, Tübingen, Germany), Javier Simón-Sánchez (Department for Neurodegenerative Diseases, Hertie Institute for Clinical Brain Research, University of Tübingen, and DZNE, German Center for Neurodegenerative Diseases, Tübingen, Germany), Peter Heutink (DZNE, German Center for Neurodegenerative Diseases and Department for Neurodegenerative Diseases, Hertie Institute for Clinical Brain Research, University of Tübingen, Tübingen, Germany), Patrizia Rizzu (DZNE, German Center for Neurodegenerative Diseases), Manu Sharma (Centre for Genetic Epidemiology, Institute for Clinical Epidemiology and Applied Biometry, University of Tubingen, Germany), Thomas Gasser (Department for Neurodegenerative Diseases, Hertie Institute for Clinical Brain Research, and DZNE, German Center for Neurodegenerative Diseases, Tübingen, Germany),

**United States of America:** Mark R Cookson (Laboratory of Neurogenetics, National Institute on Aging, Bethesda, USA), Sara Bandres-Ciga (Laboratory of Neurogenetics, National Institute on Aging, Bethesda, MD, USA), Cornelis Blauwendraat (National Institute on Aging and National Institute of Neurological Disorders and Stroke, USA), David W. Craig (Department of Translational Genomics, Keck School of Medicine, University of Southern California, Los Angeles, USA), Derek Narendra (Inherited Movement Disorders Unit, National Institute of Neurological Disorders and Stroke, Bethesda, MD, USA), Faraz Faghri (Laboratory of Neurogenetics, National Institute on Aging, Bethesda, USA; Department of Computer Science, University of Illinois at Urbana-Champaign, Urbana, IL, USA), J Raphael Gibbs (Laboratory of Neurogenetics, National Institute on Aging, National Institutes of Health, Bethesda, MD, USA), Dena G. Hernandez (Laboratory of Neurogenetics, National Institute on Aging, Bethesda, MD, USA), Kendall Van Keuren-Jensen (Neurogenomics Division, TGen, Phoenix, AZ USA), Joshua M. Shulman (Departments of Neurology, Neuroscience, and Molecular & Human Genetics, Baylor College of Medicine, Houston, Texas, USA; Jan and Dan Duncan Neurological Research Institute, Texas Children’s Hospital, Houston, Texas, USA), Hirotaka Iwaki (Laboratory of Neurogenetics, National Institute on Aging, Bethesda, MD, USA), Hampton L. Leonard (Laboratory of Neurogenetics, National Institute on Aging, Bethesda, MD, USA), Mike A. Nalls (Laboratory of Neurogenetics, National Institute on Aging, Bethesda, USA; CEO/Consultant Data Tecnica International, Glen Echo, MD, USA), Laurie Robak (Baylor College of Medicine, Houston, Texas, USA), Jose Bras (Center for Neurodegenerative Science, Van Andel Research Institute, Grand Rapids, Michigan, USA), Rita Guerreiro (Center for Neurodegenerative Science, Van Andel Research Institute, Grand Rapids, Michigan, USA), Steven Lubbe (Ken and Ruth Davee Department of Neurology, Northwestern University Feinberg School of Medicine, Chicago, IL, USA), Steven Finkbeiner (Departments of Neurology and Physiology, University of California, San Francisco; Gladstone Institute of Neurological Disease; Taube/Koret Center for Neurodegenerative Disease Research, San Francisco, CA, USA), Niccolo E. Mencacci (Northwestern University Feinberg School of Medicine, Chicago, IL, USA), Codrin Lungu (National Institutes of Health Division of Clinical Research, NINDS, National Institutes of Health, Bethesda, MD, USA), Andrew B Singleton (Laboratory of Neurogenetics, National Institute on Aging, Bethesda, MD, USA), Sonja W. Scholz (Neurodegenerative Diseases Research Unit, National Institute of Neurological Disorders and Stroke, Bethesda, MD, USA), Xylena Reed (Laboratory of Neurogenetics, National Institute on Aging, Bethesda, MD, USA). Roy N. Alcalay (Department of Neurology, College of Physicians and Surgeons, Columbia University Medical Center, New York, NY, USA, Taub Institute for Research on Alzheimer's Disease and the Aging Brain, College of Physicians and Surgeons, Columbia University Medical Center, New York, NY, USA)

**Canada:** Ziv Gan-Or (Montreal Neurological Institute and Hospital, Department of Neurology & Neurosurgery, Department of Human Genetics, McGill University, Montréal, QC, H3A 0G4, Canada), Guy A. Rouleau (Montreal Neurological Institute and Hospital, Department of Neurology & Neurosurgery, Department of Human Genetics, McGill University, Montréal, QC, H3A 0G4, Canada), Lynne Krohn (Montreal Neurological Institute and Hospital, Department of Neurology & Neurosurgery, Department of Human Genetics, McGill University, Montréal, QC, H3A 0G4, Canada), Lynne Krohn (Montreal Neurological Institute and Hospital, Department of Neurology & Neurosurgery, Department of Human Genetics, McGill University, Montréal, QC, H3A 0G4, Canada),

**The Netherlands:** Jacobus J van Hilten (Department of Neurology, Leiden University Medical Center, Leiden, Netherlands), Johan Marinus (Department of Neurology, Leiden University Medical Center, Leiden, Netherlands)

**Spain:** Astrid D. Adarmes-Gómez (Instituto de Biomedicina de Sevilla (IBiS), Hospital Universitario Virgen del Rocío/CSIC/Universidad de Sevilla, Seville), Miquel Aguilar (Fundació Docència i Recerca Mútua de Terrassa and Movement Disorders Unit, Department of Neurology, University Hospital Mutua de Terrassa, Terrassa, Barcelona.), Ignacio Alvarez (Fundació Docència i Recerca Mútua de Terrassa and Movement Disorders Unit, Department of Neurology, University Hospital Mutua de Terrassa, Terrassa, Barcelona.),Victoria Alvarez (Hospital Universitario Central de Asturias, Oviedo), Francisco Javier Barrero (Hospital Universitario San Cecilio de Granada, Universidad de Granada), Jesús Alberto Bergareche Yarza (Instituto de Investigación Sanitaria Biodonostia, San Sebastián), Inmaculada Bernal-Bernal (Instituto de Biomedicina de Sevilla (IBiS), Hospital Universitario Virgen del Rocío/CSIC/Universidad de Sevilla, Seville), Marta Blazquez (Hospital Universitario Central de Asturias, Oviedo), Marta Bonilla-Toribio (Instituto de Biomedicina de Sevilla (IBiS), Hospital Universitario Virgen del Rocío/CSIC/Universidad de Sevilla, Seville), Juan A. Botía (Universidad de Murcia, Murcia), María Teresa Boungiorno (Fundació Docència i Recerca Mútua de Terrassa and Movement Disorders Unit, Department of Neurology, University Hospital Mutua de Terrassa, Terrassa, Barcelona.) Dolores Buiza-Rueda (Instituto de Biomedicina de Sevilla (IBiS), Hospital Universitario Virgen del Rocío/CSIC/Universidad de Sevilla, Seville), Ana Cámara (Hospital Clinic de Barcelona), Fátima Carrillo (Instituto de Biomedicina de Sevilla (IBiS), Hospital Universitario Virgen del Rocío/CSIC/Universidad de Sevilla, Seville), Mario Carrión-Claro (Instituto de Biomedicina de Sevilla (IBiS), Hospital Universitario Virgen del Rocío/CSIC/Universidad de Sevilla, Seville), Debora Cerdan (Hospital General de Segovia, Segovia), Jordi Clarimón (Memory Unit, Department of Neurology, IIB Sant Pau, Hospital de la Santa Creu i Sant Pau, Universitat Autònoma de Barcelona and Centro de Investigación Biomédica en Red en Enfermedades Neurodegenerativas (CIBERNED), Madrid),Yaroslau Compta (Hospital Clinic de Barcelona), Monica Diez-Fairen (Fundació Docència i Recerca Mútua de Terrassa and Movement Disorders Unit, Department of Neurology, University Hospital Mutua de Terrassa, Terrassa, Barcelona.), Oriol Dols-Icardo (Memory Unit, Department of Neurology, IIB Sant Pau, Hospital de la Santa Creu i Sant Pau, Universitat Autònoma de Barcelona, Barcelona, and Centro de Investigación Biomédica en Red en Enfermedades Neurodegenerativas (CIBERNED), Madrid), Jacinto Duarte (Hospital General de Segovia, Segovia), Raquel Duran (Centro de Investigacion Biomedica, Universidad de Granada, Granada), Francisco Escamilla-Sevilla (Hospital Universitario Virgen de las Nieves, Instituto de Investigación Biosanitaria de Granada, Granada), Mario Ezquerra (Hospital Clinic de Barcelona), Cici Feliz (Departmento de Neurologia, Instituto de Investigación Sanitaria Fundación Jiménez Díaz, Madrid, Spain), Manel Fernández (Hospital Clinic de Barcelona), Rubén Fernández-Santiago (Hospital Clinic de Barcelona), Ciara Garcia (Hospital Universitario Central de Asturias, Oviedo), Pedro García-Ruiz (Instituto de Investigación Sanitaria Fundación Jiménez Díaz, Madrid), Pilar Gómez-Garre (Instituto de Biomedicina de Sevilla (IBiS), Hospital Universitario Virgen del Rocío/CSIC/Universidad de Sevilla, Seville), Maria Jose Gomez Heredia (Hospital Universitario Virgen de la Victoria, Malaga), Isabel Gonzalez-Aramburu (Hospital Universitario Marqués de Valdecilla-IDIVAL, Santander) ,Ana Gorostidi Pagola (Instituto de Investigación Sanitaria Biodonostia, San Sebastián), Janet Hoenicka (Institut de Recerca Sant Joan de Déu, Barcelona), Jon Infante (Hospital Universitario Marqués de Valdecilla-IDIVAL and University of Cantabria, Santander, and Centro de Investigación Biomédica en Red en Enfermedades Neurodegenerativas (CIBERNED), Silvia Jesús (Instituto de Biomedicina de Sevilla (IBiS), Hospital Universitario Virgen del Rocío/CSIC/Universidad de Sevilla, Seville), Adriano Jimenez-Escrig (Hospital Universitario Ramón y Cajal, Madrid), Jaime Kulisevsky (Movement Disorders Unit, Department of Neurology, IIB Sant Pau, Hospital de la Santa Creu i Sant Pau, Universitat Autònoma de Barcelona, Barcelona, and Centro de Investigación Biomédica en Red en Enfermedades Neurodegenerativas (CIBERNED)), Miguel A. Labrador-Espinosa (Instituto de Biomedicina de Sevilla (IBiS), Hospital Universitario Virgen del Rocío/CSIC/Universidad de Sevilla, Seville), Jose Luis Lopez-Sendon (Hospital Universitario Ramón y Cajal, Madrid), Adolfo López de Munain Arregui (Instituto de Investigación Sanitaria Biodonostia, San Sebastián), Daniel Macias (Instituto de Biomedicina de Sevilla (IBiS), Hospital Universitario Virgen del Rocío/CSIC/Universidad de Sevilla, Seville), Irene Martínez Torres (Department of Neurology, Instituto de Investigación Sanitaria La Fe, Hospital Universitario y Politécnico La Fe, Valencia), Juan Marín (Movement Disorders Unit, Department of Neurology, IIB Sant Pau, Hospital de la Santa Creu i Sant Pau, Universitat Autònoma de Barcelona, Barcelona, and Centro de Investigación Biomédica en Red en Enfermedades Neurodegenerativas (CIBERNED)), Maria Jose Marti (Hospital Clinic Barcelona), Juan Carlos Martínez-Castrillo (Instituto Ramón y Cajal de Investigación Sanitaria, Hospital Universitario Ramón y Cajal, Madrid), Carlota Méndez-del-Barrio (Instituto de Biomedicina de Sevilla (IBiS), Hospital Universitario Virgen del Rocío/CSIC/Universidad de Sevilla, Seville), Manuel Menéndez González (Hospital Universitario Central de Asturias, Oviedo), Marina Mata (Department of Neurology, Hospital Universitario Infanta Sofía, Madrid, Spain) Adolfo Mínguez (Hospital Universitario Virgen de las Nieves, Granada, Instituto de Investigación Biosanitaria de Granada), Pablo Mir (Instituto de Biomedicina de Sevilla (IBiS), Hospital Universitario Virgen del Rocío/CSIC/Universidad de Sevilla, Seville), Elisabet Mondragon Rezola (Instituto de Investigación Sanitaria Biodonostia, San Sebastián), Esteban Muñoz (Hospital Clinic Barcelona), Javier Pagonabarraga (Movement Disorders Unit, Department of Neurology, IIB Sant Pau, Hospital de la Santa Creu i Sant Pau, Universitat Autònoma de Barcelona, Barcelona, and Centro de Investigación Biomédica en Red en Enfermedades Neurodegenerativas (CIBERNED)), Pau Pastor (Fundació Docència i Recerca Mútua de Terrassa and Movement Disorders Unit, Department of Neurology, University Hospital Mutua de Terrassa, Terrassa, Barcelona.), Francisco Perez Errazquin (Hospital Universitario Virgen de la Victoria, Malaga), Teresa Periñán-Tocino (Instituto de Biomedicina de Sevilla (IBiS), Hospital Universitario Virgen del Rocío/CSIC/Universidad de Sevilla, Seville), Javier Ruiz-Martínez (Hospital Universitario Donostia, Instituto de Investigación Sanitaria Biodonostia, San Sebastián), Clara Ruz (Centro de Investigacion Biomedica, Universidad de Granada, Granada), Antonio Sanchez Rodriguez (Hospital Universitario Marqués de Valdecilla-IDIVAL, Santander), María Sierra (Hospital Universitario Marqués de Valdecilla-IDIVAL, Santander), Esther Suarez-Sanmartin (Hospital Universitario Central de Asturias, Oviedo), Cesar Tabernero (Hospital General de Segovia, Segovia), Juan Pablo Tartari (Fundació Docència i Recerca Mútua de Terrassa and Movement Disorders Unit, Department of Neurology, University Hospital Mutua de Terrassa, Terrassa, Barcelona), Cristina Tejera-Parrado (Instituto de Biomedicina de Sevilla (IBiS), Hospital Universitario Virgen del Rocío/CSIC/Universidad de Sevilla, Seville), Eduard Tolosa (Hospital Clinic Barcelona), Francesc Valldeoriola (Hospital Clinic Barcelona), Laura Vargas-González (Instituto de Biomedicina de Sevilla (IBiS), Hospital Universitario Virgen del Rocío/CSIC/Universidad de Sevilla, Seville), Lydia Vela (Department of Neurology, Hospital Universitario Fundación Alcorcón, Madrid), Francisco Vives (Centro de Investigacion Biomedica, Universidad de Granada, Granada).

**Austria:** Alexander Zimprich (Department of Neurology, Medical University of Vienna, Austria)

**Norway:** Lasse Pihlstrom (Department of Neurology, Oslo University Hospital, Oslo, Norway), Mathias Toft (Department of Neurology and Institute of Clinical Medicine, Oslo University Hospital, Oslo, Norway)

**Estonia**: Sulev Koks (Department of Pathophysiology, University of Tartu, Tartu, Estonia; Department of Reproductive Biology, Estonian University of Life Sciences, Tartu, Estonia), Pille Taba (Department of Neurology and Neurosurgery, University of Tartu, Tartu, Estonia)

**Israel:** Sharon Hassin-Baer (The Movement Disorders Institute, Department of Neurology and Sagol Neuroscience Center, Chaim Sheba Medical Center, Tel-Hashomer, 5262101, Ramat Gan, Israel, Sackler Faculty of Medicine, Tel Aviv University, Tel Aviv, Israel)

**IPDGC consortium acknowledgement:**

We would like to thank all of the subjects who donated their time and biological samples to be a part of this study. We also would like to thank all members of the International Parkinson Disease Genomics Consortium (IPDGC). See for a complete overview of members, acknowledgements and funding http://pdgenetics.org/partners.This work was supported in part by the Intramural Research Programs of the National Institute of Neurological Disorders and Stroke (NINDS), the National Institute on Aging (NIA), and the National Institute of Environmental Health Sciences both part of the National Institutes of Health, Department of Health and Human Services; project numbers 1ZIA-NS003154, Z01-AG000949-02 and Z01-ES101986. In addition this work was supported by the Department of Defense (award W81XWH-09-2-0128), and The Michael J Fox Foundation for Parkinson’s Research. This work was supported by National Institutes of Health grants R01NS037167, R01CA141668, P50NS071674, American Parkinson Disease Association (APDA); Barnes Jewish Hospital Foundation; Greater St Louis Chapter of the APDA. The KORA (Cooperative Research in the Region of Augsburg) research platform was started and financed by the Forschungszentrum für Umwelt und Gesundheit, which is funded by the German Federal Ministry of Education, Science, Research, and Technology and by the State of Bavaria. This study was also funded by the German Federal Ministry of Education and Research (BMBF) under the funding code 031A430A, the EU Joint Programme - Neurodegenerative Diseases Research (JPND) project under the aegis of JPND -www.jpnd.eu- through Germany, BMBF, funding code 01ED1406 and iMed - the Helmholtz Initiative on Personalized Medicine. This study is funded by the German National Foundation grant (DFG SH599/6-1) (grant to M.S), Michael J Fox Foundation, and MSA Coalition, USA (to M.S). The French GWAS work was supported by the French National Agency of Research (ANR-08-MNP-012). This study was also funded by France-Parkinson Association, Fondation de France, the French program “Investissements d’avenir” funding (ANR-10-IAIHU-06) and a grant from Assistance Publique-Hôpitaux de Paris (PHRC, AOR-08010) for the French clinical data. This study was also sponsored by the Landspitali University Hospital Research Fund (grant to SSv); Icelandic Research Council (grant to SSv); and European Community Framework Programme 7, People Programme, and IAPP on novel genetic and phenotypic markers of Parkinson’s disease and Essential Tremor (MarkMD), contract number PIAP-GA-2008-230596 MarkMD (to HP and JHu). Institutional research funding IUT20-46 was received of the Estonian Ministry of Education and Research (SK). The McGill study was funded by the Michael J. Fox Foundation and the Canadian Consortium on Neurodegeneration in Aging (CCNA). This study utilized the high-performance computational capabilities of the Biowulf Linux cluster at the National Institutes of Health, Bethesda, Md. (http://biowulf.nih.gov), and DNA panels, samples, and clinical data from the National Institute of Neurological Disorders and Stroke Human Genetics Resource Center DNA and Cell Line Repository. People who contributed samples are acknowledged in descriptions of every panel on the repository website. We thank the French Parkinson’s Disease Genetics Study Group and the Drug Interaction with genes (DIGPD) study group: Y Agid, M Anheim, F Artaud, A-M Bonnet, C Bonnet, F Bourdain, J-P Brandel, C Brefel-Courbon, M Borg, A Brice, E Broussolle, F Cormier-Dequaire, J-C Corvol, P Damier, B Debilly, B Degos, P Derkinderen, A Destée, A Dürr, F Durif, A Elbaz, D Grabli, A Hartmann, S Klebe, P. Krack, J Kraemmer, S Leder, S Lesage, R Levy, E Lohmann, L Lacomblez, G Mangone, L-L Mariani, A-R Marques, M Martinez, V Mesnage, J Muellner, F Ory-Magne, F Pico, V Planté-Bordeneuve, P Pollak, O Rascol, K Tahiri, F Tison, C Tranchant, E Roze, M Tir, M Vérin, F Viallet, M Vidailhet, A You. We also thank the members of the French 3C Consortium: A Alpérovitch, C Berr, C Tzourio, and P Amouyel for allowing us to use part of the 3C cohort, and D Zelenika for support in generating the genome-wide molecular data. We thank P Tienari (Molecular Neurology Programme, Biomedicum, University of Helsinki), T Peuralinna (Department of Neurology, Helsinki University Central Hospital), L Myllykangas (Folkhalsan Institute of Genetics and Department of Pathology, University of Helsinki), and R Sulkava (Department of Public Health and General Practice Division of Geriatrics, University of Eastern Finland) for the Finnish controls (Vantaa85+ GWAS data). We used genome-wide association data generated by the Wellcome Trust Case-Control Consortium 2 (WTCCC2) from UK patients with Parkinson’s disease and UK control individuals from the 1958 Birth Cohort and National Blood Service. Genotyping of UK replication cases on ImmunoChip was part of the WTCCC2 project, which was funded by the Wellcome Trust (083948/Z/07/Z). UK population control data was made available through WTCCC1. This study was supported by the Medical Research Council and Wellcome Trust disease centre (grant WT089698/Z/09/Z to NW, JHa, and ASc). As with previous IPDGC efforts, this study makes use of data generated by the Wellcome Trust Case-Control Consortium. A full list of the investigators who contributed to the generation of the data is available from www.wtccc.org.uk. Funding for the project was provided by the Wellcome Trust under award 076113, 085475 and 090355. This study was also supported by Parkinson’s UK (grants 8047 and J-0804) and the Medical Research Council (G0700943 and G1100643). Sequencing and genotyping done in McGill University was supported by grants from the Michael J. Fox Foundation, the Canadian Consortium on Neurodegeneration in Aging (CCNA), the Canada First Research Excellence Fund (CFREF), awarded to McGill University for the Healthy Brains for Healthy Lives (HBHL) program and Parkinson’s Society Canada. We thank Jeffrey Barrett and Jason Downing (Illumina Inc) for assistance with the design of the ImmunoChip and NeuroX arrays. DNA extraction work that was done in the UK was undertaken at University College London Hospitals, University College London, who received a proportion of funding from the Department of Health’s National Institute for Health Research Biomedical Research Centres funding. This study was supported in part by the Wellcome Trust/Medical Research Council Joint Call in Neurodegeneration award (WT089698) to the Parkinson’s Disease Consortium (UKPDC), whose members are from the UCL Institute of Neurology, University of Sheffield, and the Medical Research Council Protein Phosphorylation Unit at the University of Dundee. We thank the Quebec Parkinson’s Network (http://rpq-qpn.org) and its members. This work was supported by the Medical Research Council grant MR/N026004/1. The Braineac project was supported by the MRC through the MRC Sudden Death Brain Bank Grant (MR/G0901254) to J.H. P.A.L. was supported by the MRC (grants MR/N026004/1 and MR/L010933/1) and Michael J. Fox Foundation for Parkinson’s Research. Mike A. Nalls’ participation is supported by a consulting contract between Data Tecnica International and the National Institute on Aging, NIH, Bethesda, MD, USA, as a possible conflict of interest Dr. Nalls also consults for Illumina Inc, Lysosomal Therapeutics Inc, the Michael J. Fox Foundation and Vivid Genomics among others.
