## Supplement for "Genetic modifiers of risk and age at onset in *GBA* associated Parkinson’s disease and Lewy body dementia": GBA modifiers Supplemental Figures final version.docx

Blauwendraat et al. 2019

***Supplemental Figures***

Supplemental Figure 1: Genetic risk score differences between GBA positive controls and GBA positive cases separated by variant groups

Supplemental Figure 2: Area under the curve including the 95% confidence interval for ROC curve plot of predictive value of Parkinson’s disease genetic risk score in GBA carriers from the IPDGC dataset

Supplemental Figure 3: Area under the curve including the 95% confidence interval for ROC curve plot of predictive value of Parkinson’s disease genetic risk score in GBA carriers from the UK Biobank dataset

Supplemental Figure 4: Area under the curve including the 95% confidence interval for ROC curve plot of predictive value of Parkinson’s disease genetic risk score in GBA carriers from the NDRU-LBD dataset

Supplemental Figure 5: Manhattan plot of GBA carriers case-control GWAS results meta-analyzed

Supplemental Figure 6: Forest plot of (PD) case control associations of *SNCA* rs356219 in the IPDGC data and UK Biobank *GBA* carriers.

Supplemental Figure 7: Forest plot of (PD) case control associations of *CTSB* rs1293298 in the IPDGC data and UK Biobank *GBA* carriers.

Supplemental Figure 8: Beta value (effect size) correlation plot between PD GWAS (Nalls et al 2019) and the GBA carrier case-control GWAS.

Supplemental Figure 9: Forest plot of the meta-analyses of the interaction between the SNCA variants rs356219 and the GBA carrier status.

Supplemental Figure 10: Expression quantitative trait loci analyses of CTSB and rs1293298 in the NABEC dataset.

Supplemental Figure 11: Expression of SNCA, GBA and CTSB in single-nuclei RNAseq data derived from three frozen prefrontal cortex samples and four frozen hippocampal samples.

Supplemental Figure 12: Immunofluorescent images of differentiated iPS cells showing expression of neuronal markers Beta-III Tubulin and MAP2.

Supplemental Figure 13: A) Western blot of α-synuclein expression in GBA WT and N370S forebrain neurons. B) Quantitation relative to TUBB3 shows no difference in expression between genotypes.

Supplemental Figure 14: Forest plot of the association between Parkinson’s disease genetic risk score and age of onset of GBA p.N370S PD cases

**Supplemental Figure 1**

Genetic risk score differences between *GBA* positive controls and *GBA* positive cases separated by variant groups: p.E326K, p.T369M and p.N370S when excluding the *GBA* region and *LRRK2* p.G2019S from the calculations. In each variant, a highly similar effect is seen clearly separating the cases and controls. IPDGC p.E326K P=2.48E-7, OR (95%CI) 1.452 (1.31-1.59); IPDGC p.T369M P=3.07E-5, OR (95%CI) 1.553 (1.34-1.76); IPDGC p.N370S P=0.0856, OR (95%CI) 1.310 (1-1.62).

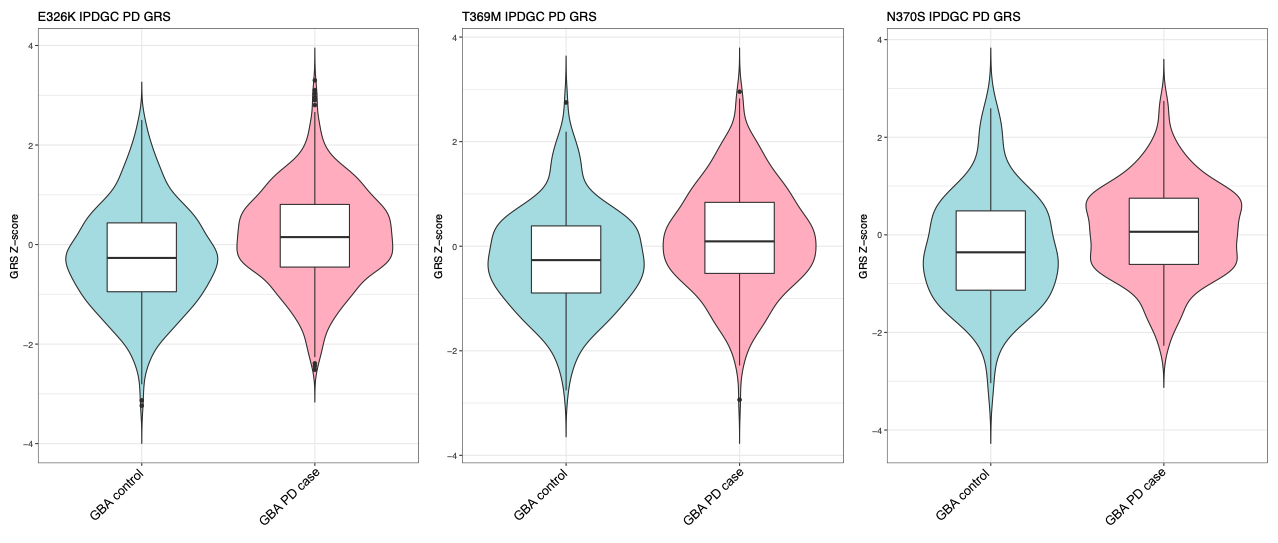

**Supplemental Figure 2**

Area under the curve including the 95% confidence interval for ROC curve plot of predictive value of Parkinson’s disease genetic risk score in GBA carriers from the IPDGC dataset

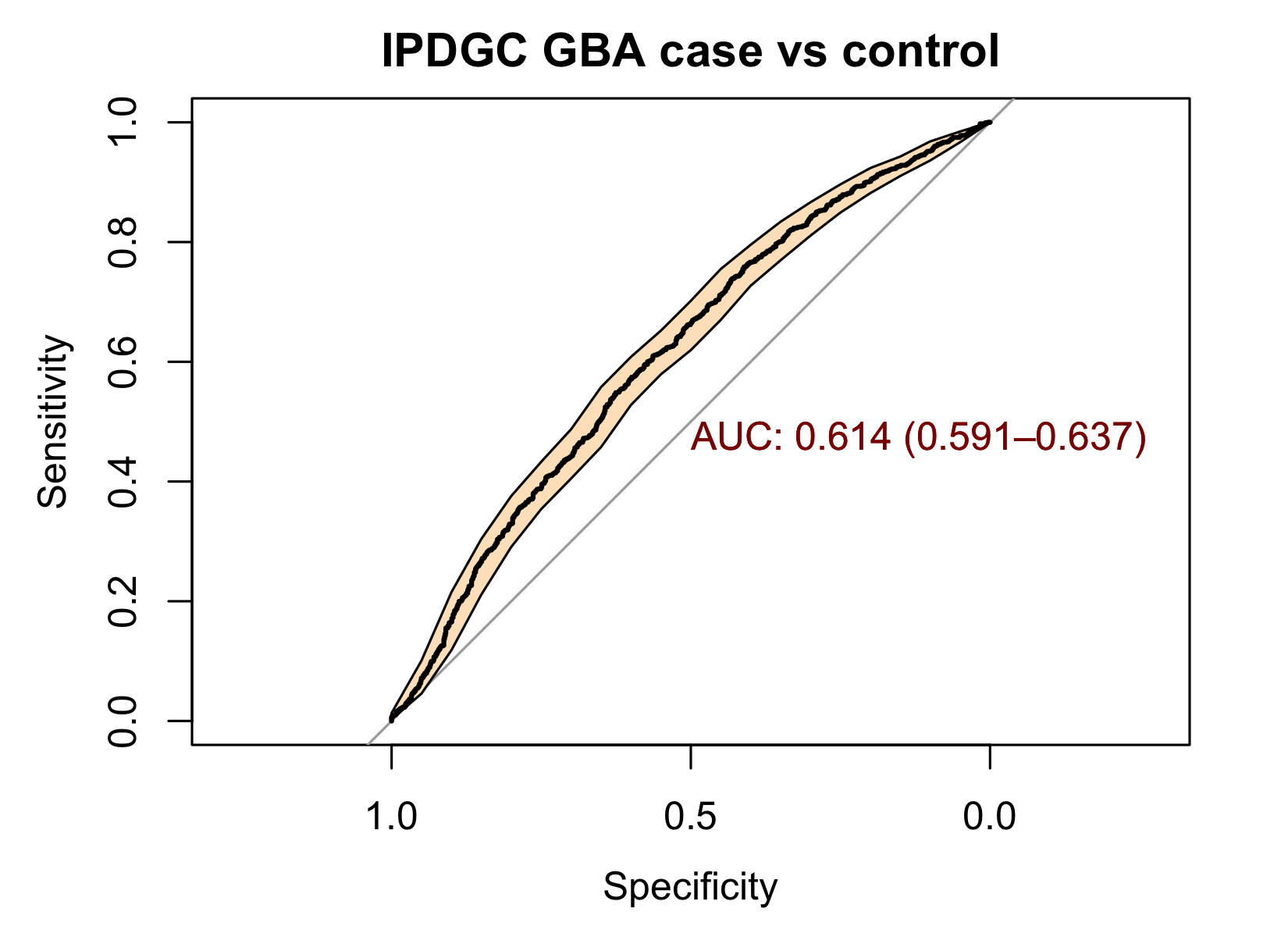

**Supplemental Figure 3**

Area under the curve including the 95% confidence interval for ROC curve plot of predictive value of Parkinson’s disease genetic risk score in GBA carriers from the UK Biobank dataset

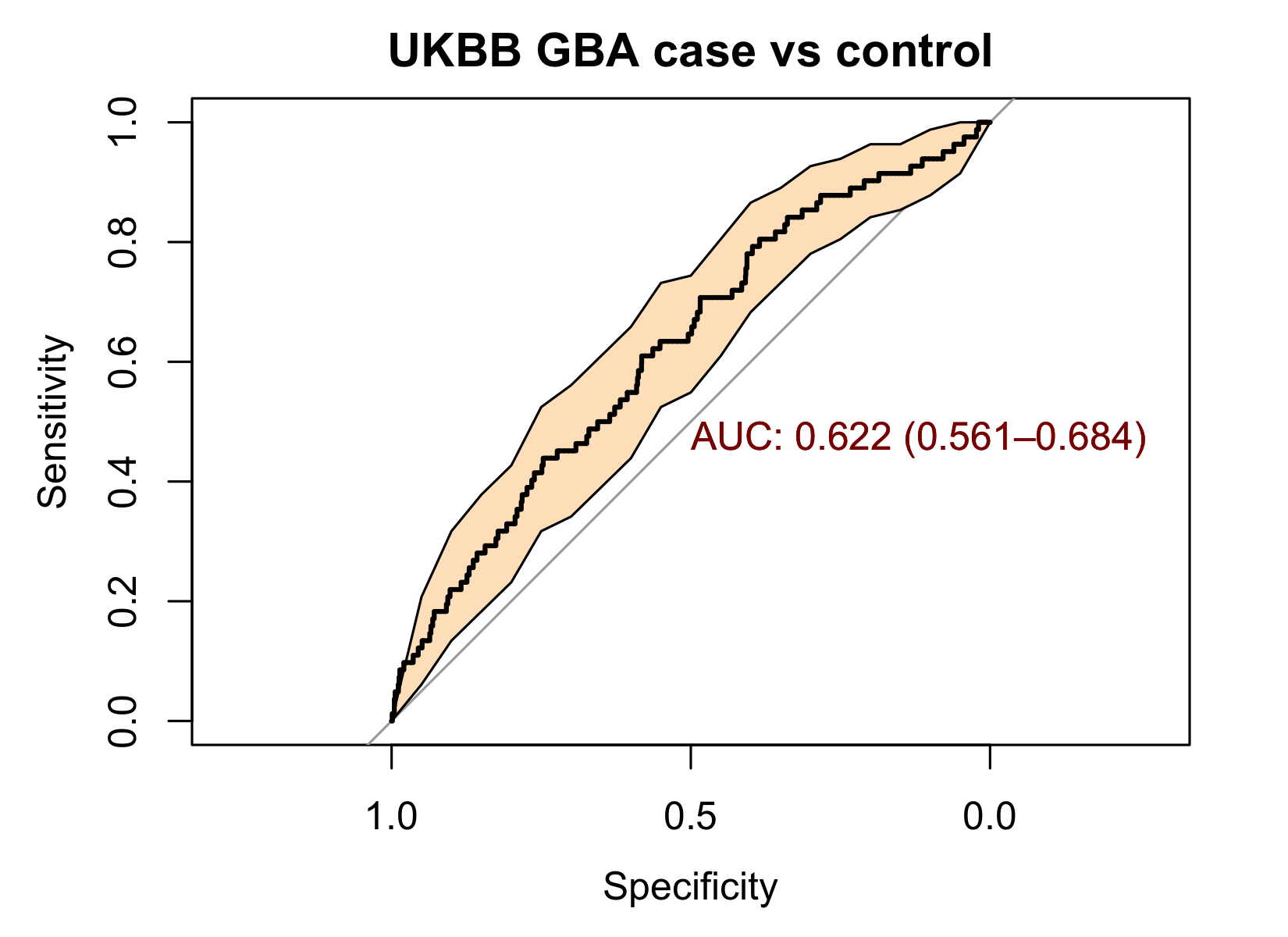

**Supplemental Figure 4**

Area under the curve including the 95% confidence interval for ROC curve plot of predictive value of Parkinson’s disease genetic risk score in GBA carriers from the NDRU-LBD dataset

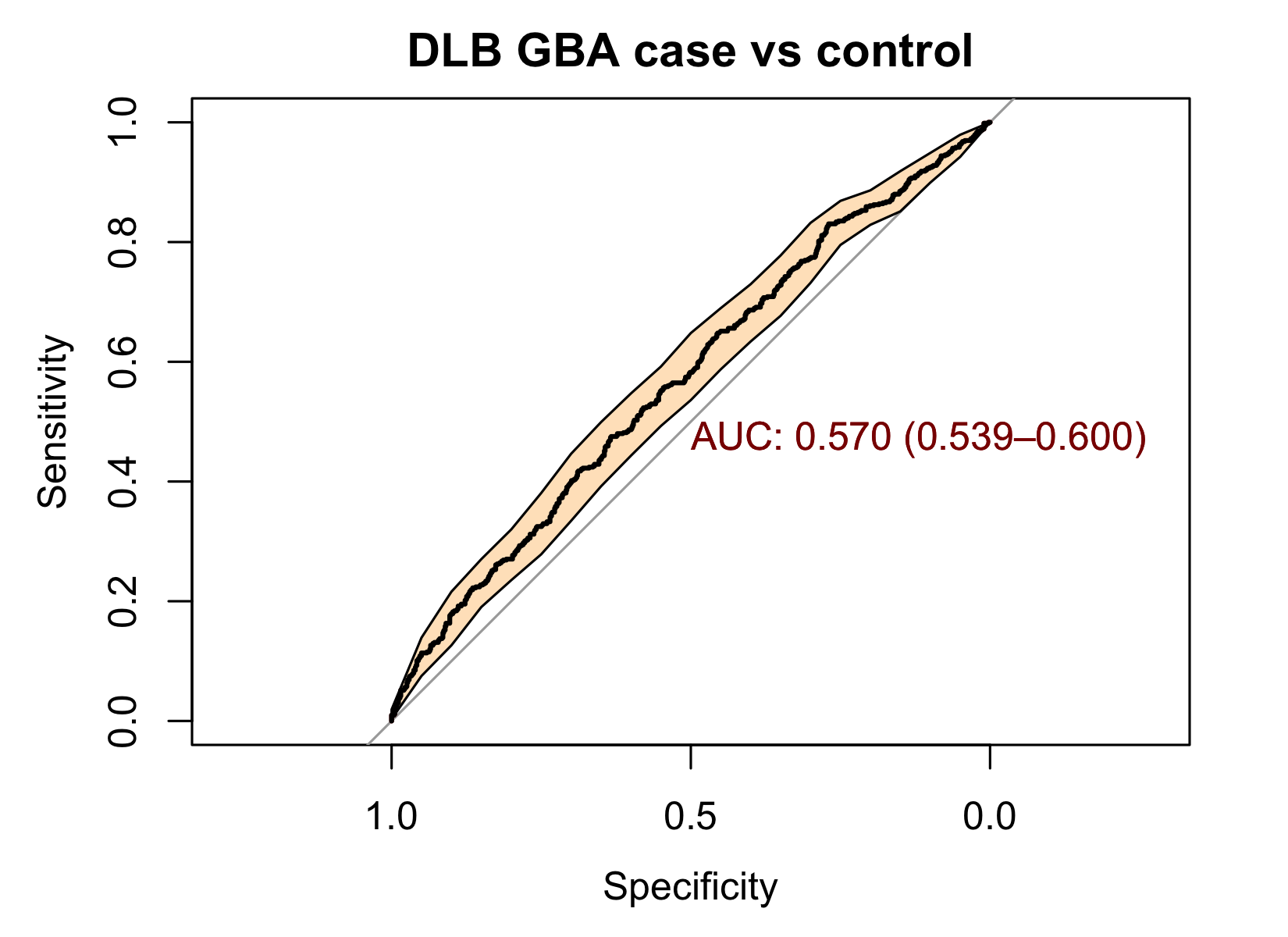

**Supplemental Figure 5**

Manhattan plot of *GBA* carriers case-control GWAS results meta-analyzed from the IPDGC and UK Biobank cohorts.

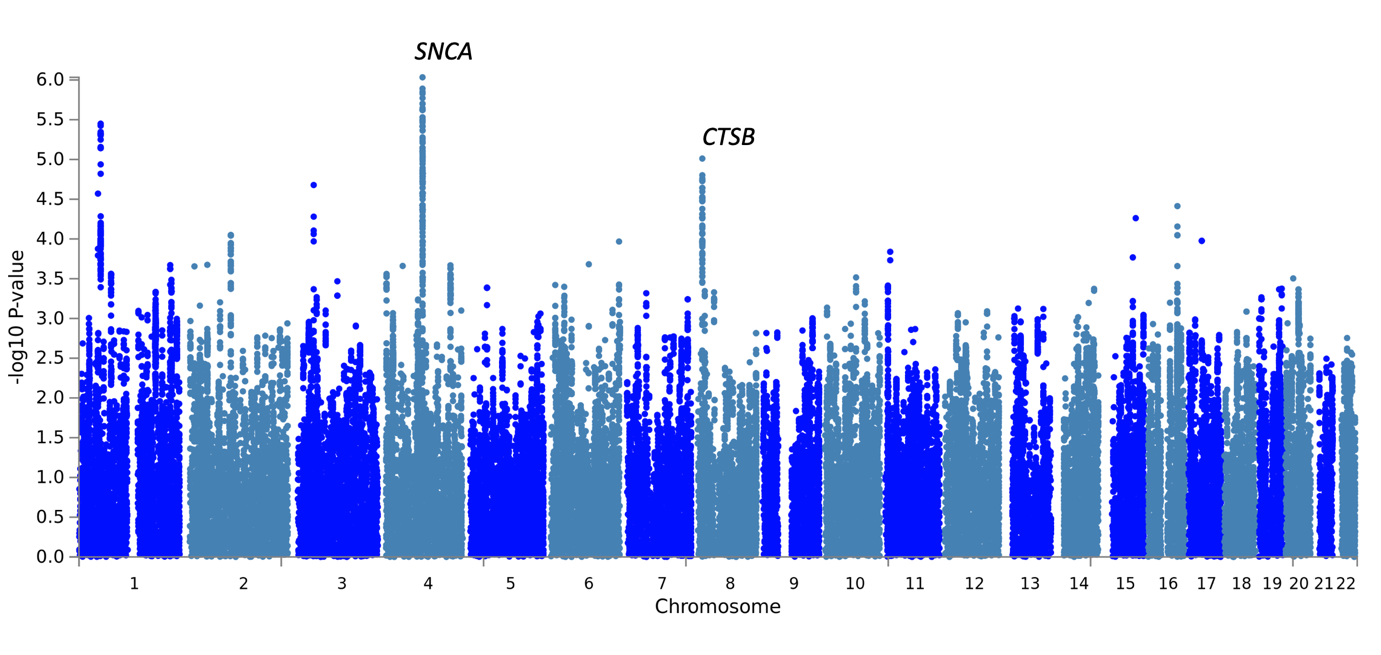

**Supplemental Figure 6**

Forest plot of (PD) case control associations of *SNCA* rs356219 in the IPDGC data and UK Biobank *GBA* carriers. On the left all *GBA* carriers are merged in the IPDGC data and on the right *GBA* carriers are separated by *GBA* variant all showing a consistent signal. Note that the LBD had a weaker effect size, but in the same direction (OR=1.112).

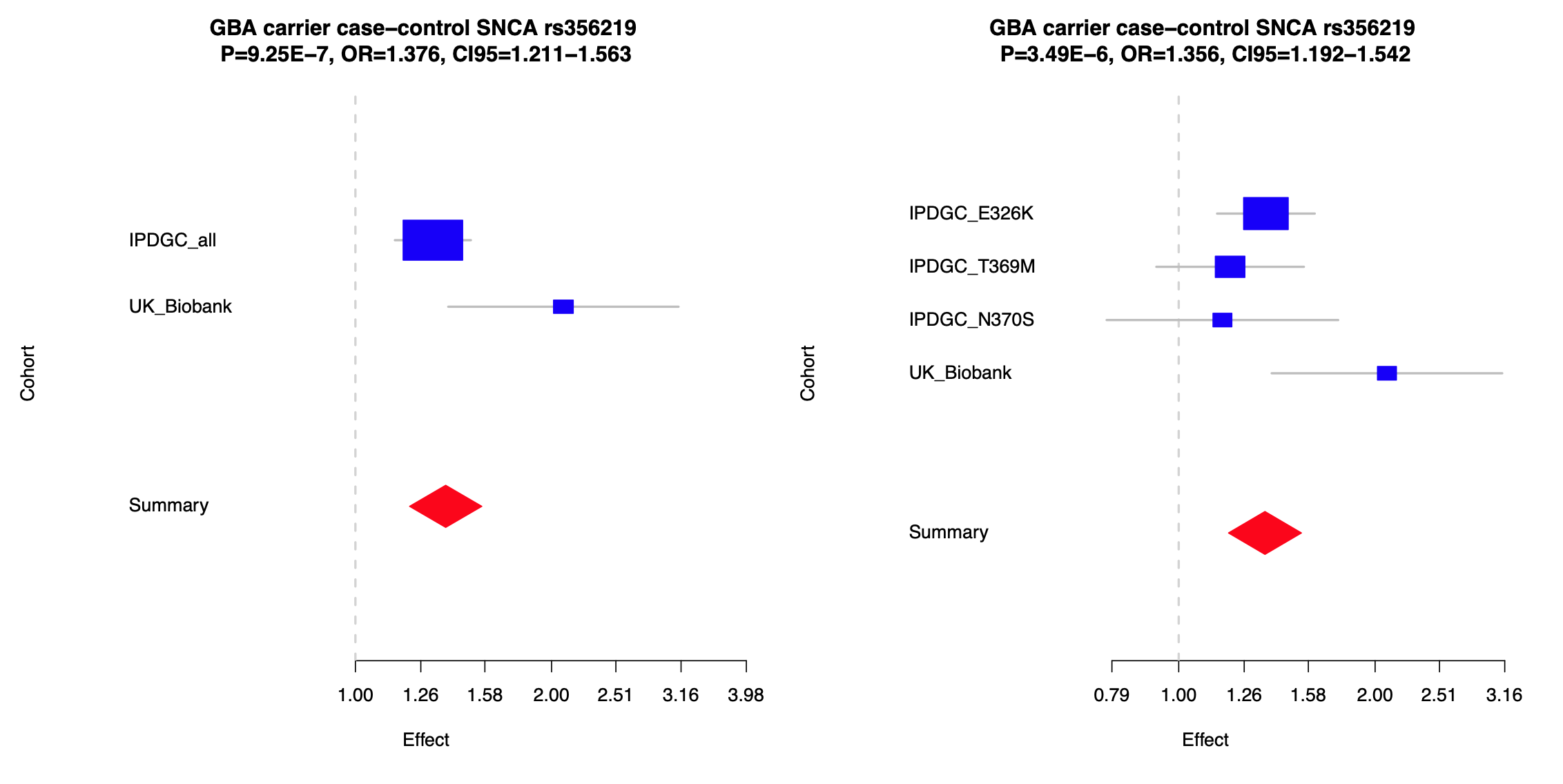

**Supplemental Figure 7**

Forest plot of (PD) case control associations of *CTSB* rs1293298 in the IPDGC data and UK Biobank *GBA* carriers. On the left all *GBA* carriers are merged in the IPDGC data and on the right *GBA* carriers are separated by *GBA* variant all showing a consistent signal. Note that the LBD had a very similar effect size (OR=0.770).

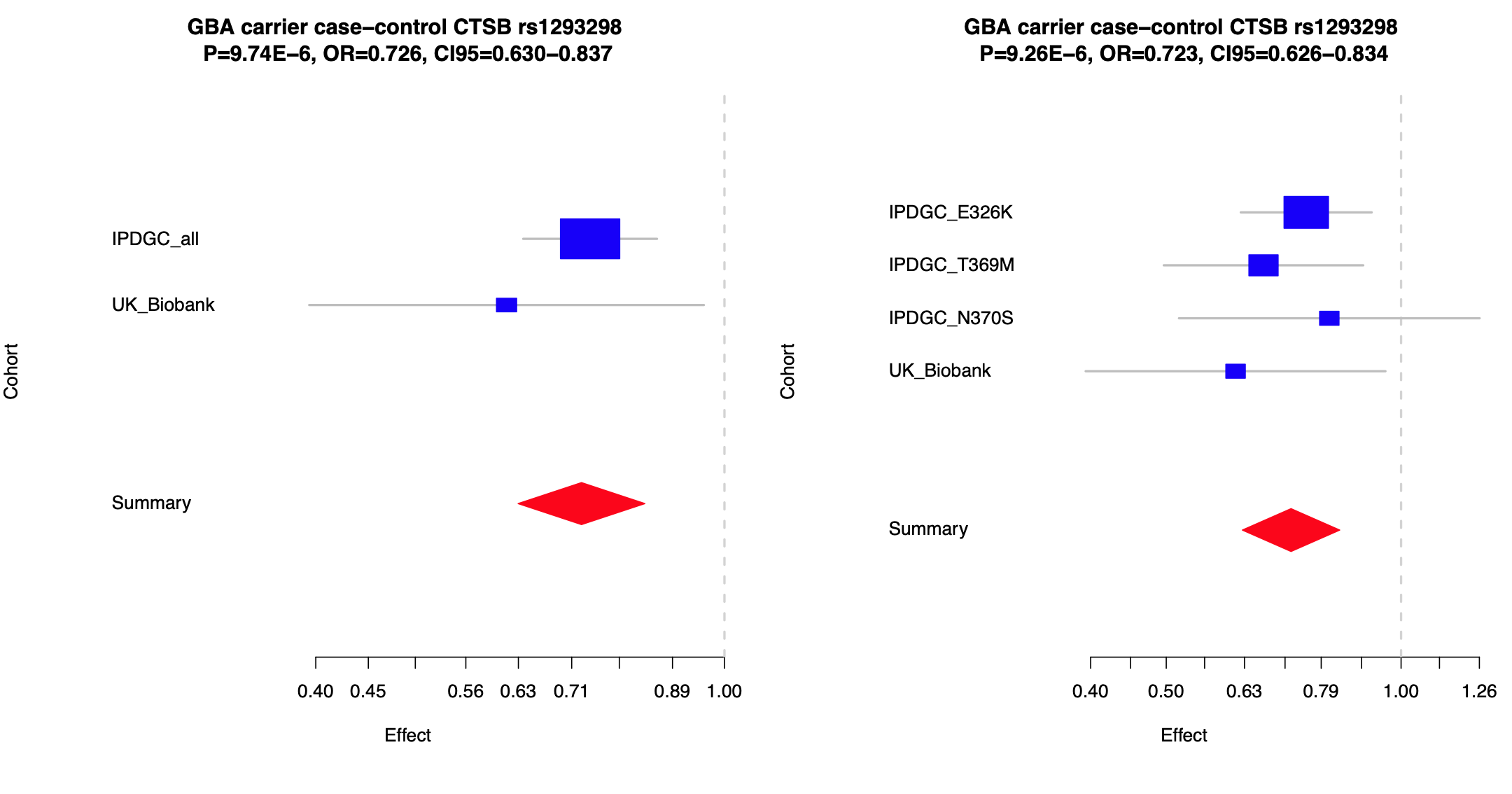

**Supplemental Figure 8**

Beta value (effect size) correlation plot between PD GWAS (Nalls et al 2018) and the GBA carrier case-control GWAS. Beta-values were plotted of the Nalls et al 2018 PD GWAS (x-axis) and the current GBA carrier case-control GWAS (y-axis). Green dots are loci that pass Bonferroni correction and red are loci that did not pass Bonferroni correction. Bonferroni correction passing SNPs are annotated by their closest gene.

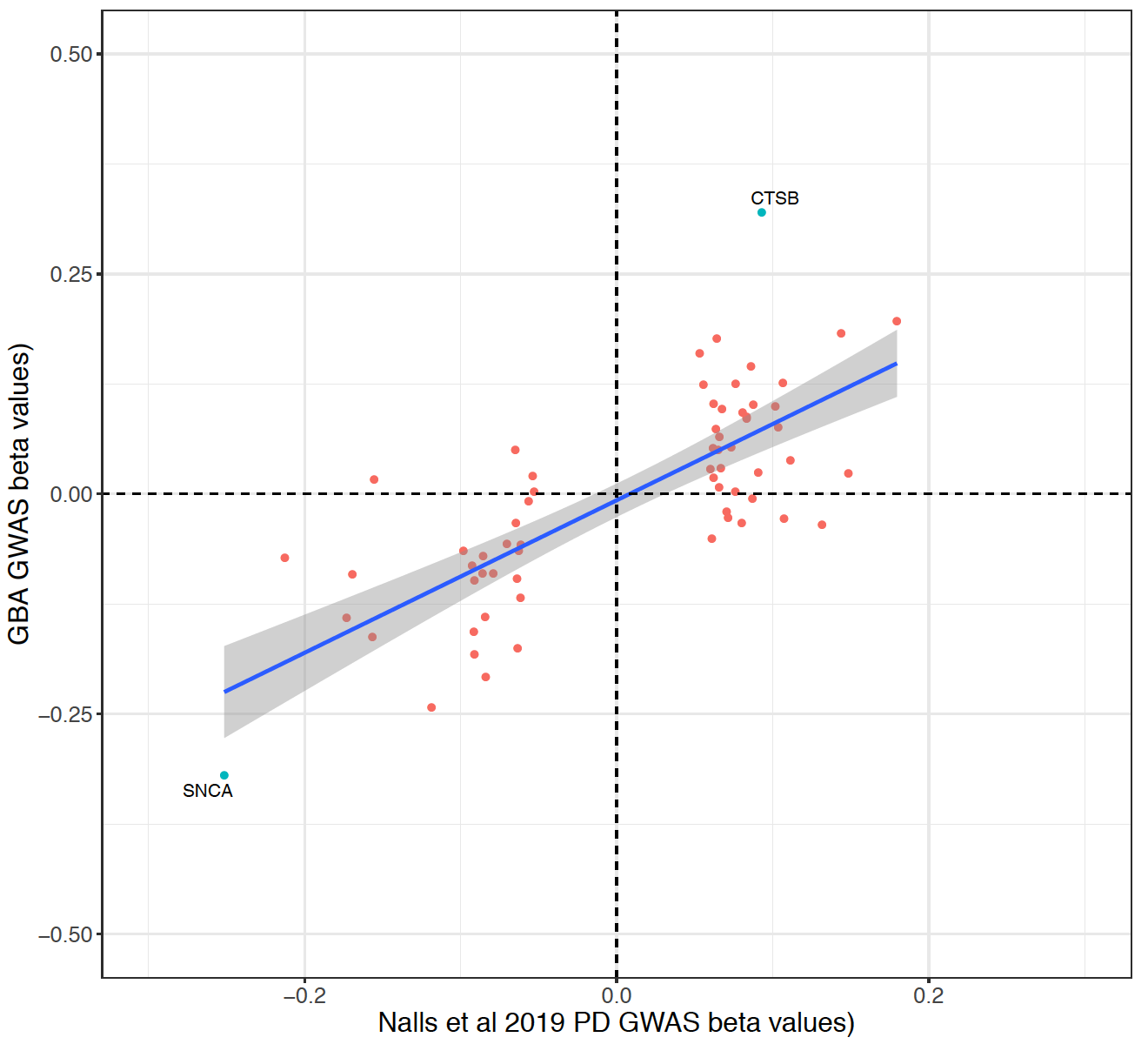

**Supplemental Figure 9**

Forest plot of the meta-analyses of the interaction between the SNCA variants rs356219 and the GBA carrier status.

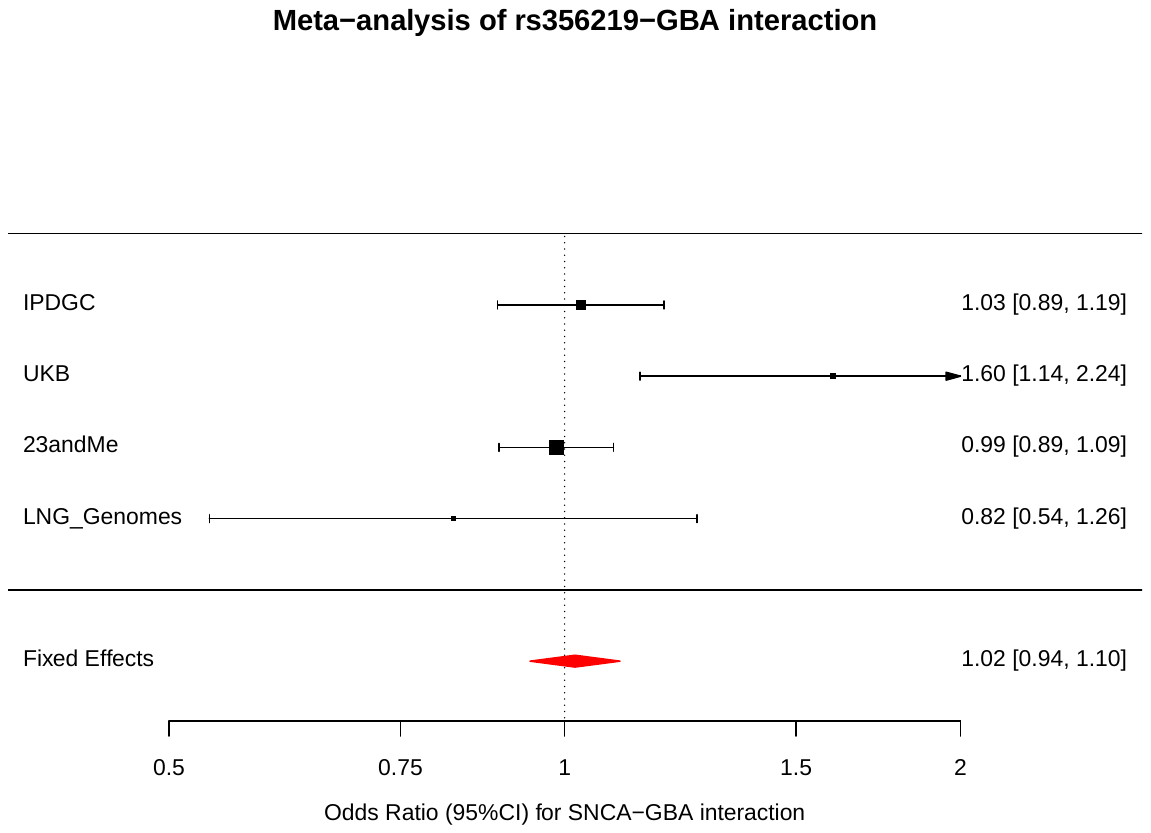

**Supplemental Figure 10**

Expression quantitative trait loci analyses of CTSB and rs1293298 in the NABEC dataset. Four transcripts were identified and the main expressed transcript uc003wuq was the only significant expression quantitative trait locus.

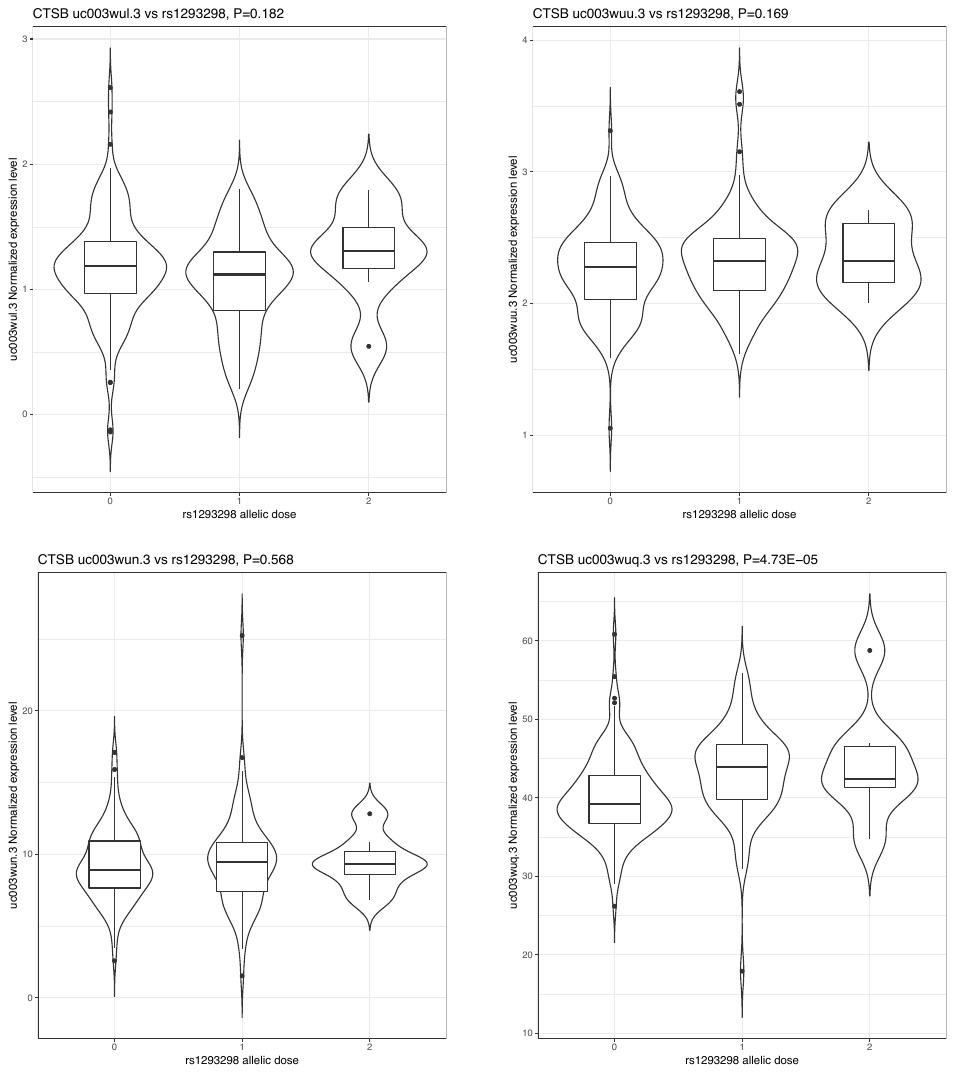

**Supplemental Figure 11**

Expression of SNCA, GBA and CTSB in single-nuclei RNAseq data derived from three frozen prefrontal cortex samples and four frozen hippocampal samples. Data was obtained from dbGaP accession phs000424.v8.p1 (Habib, Avraham-Davidi et al. 2017).

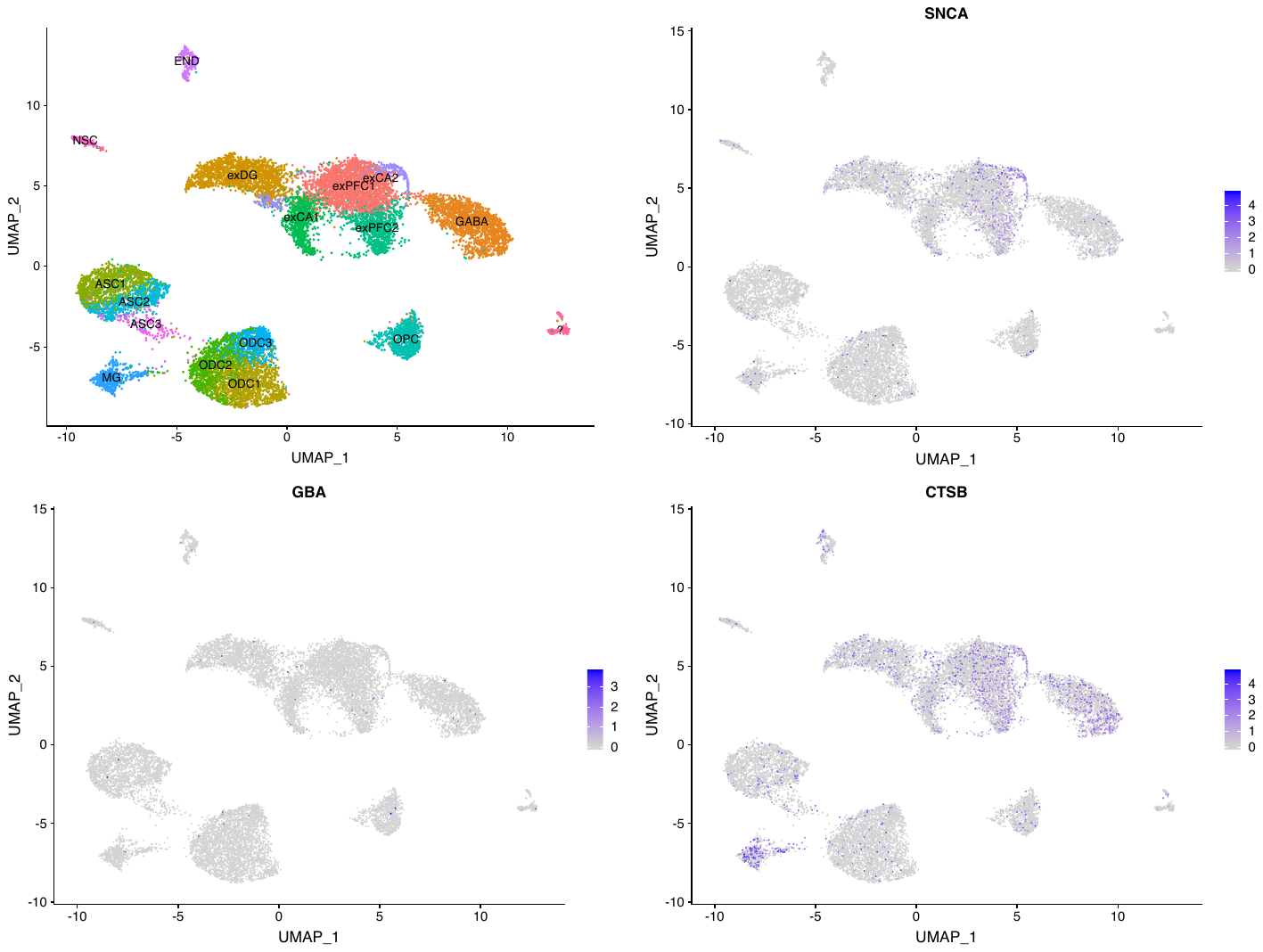

**Supplemental Figure 19:**

Immunofluorescent images of differentiated iPS cells showing expression of neuronal markers Beta-III Tubulin and MAP2.

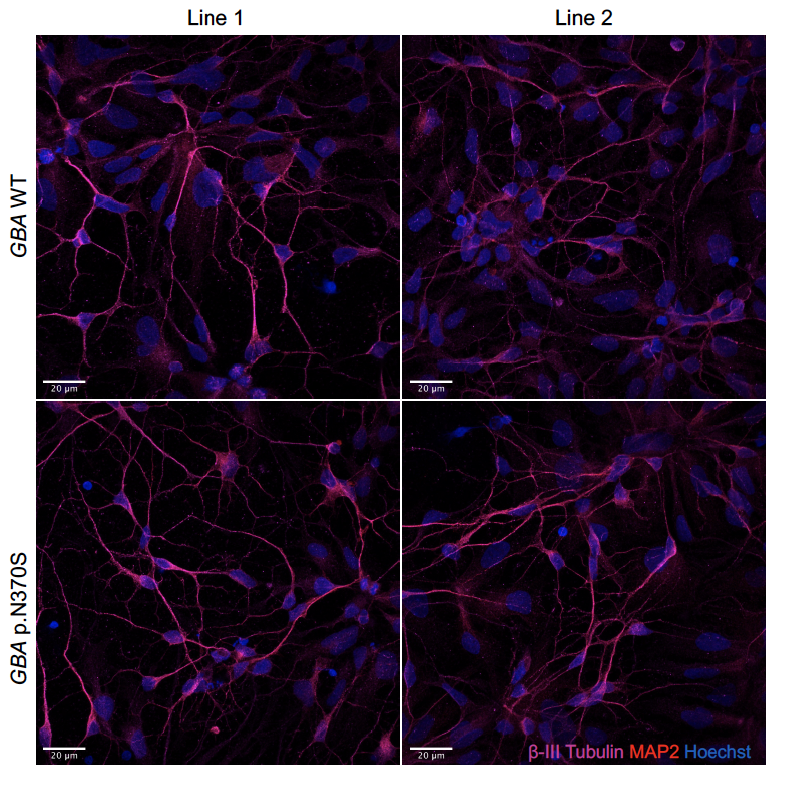

**Supplemental Figure 20**

A) Western blot of α-synuclein expression in GBA WT and N370S forebrain neurons. B) Quantitation relative to TUBB3 shows no difference in expression between genotypes.

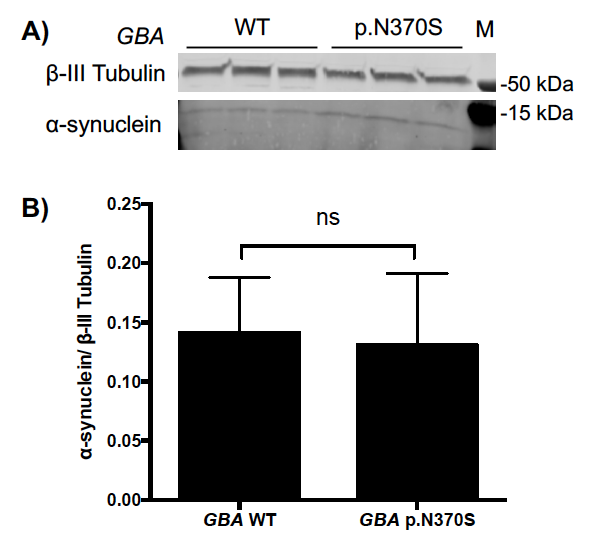

**Supplemental Figure 14**

Forest plot of the association between Parkinson’s disease genetic risk score and age of onset of GBA p.N370S PD cases.

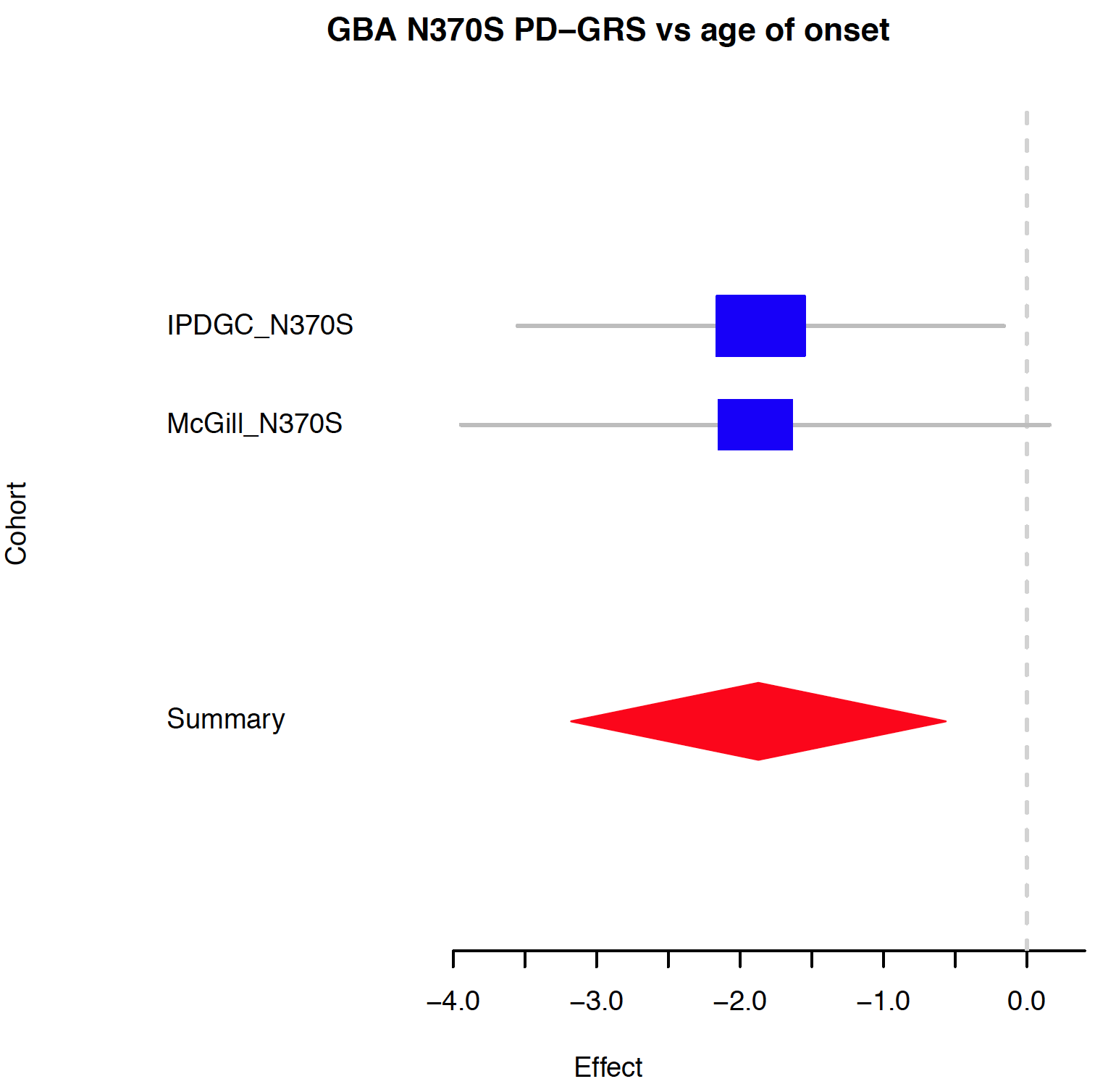
